# Engineering Spatial Control of Bacterial Organelles

**DOI:** 10.1101/2025.09.22.677801

**Authors:** Y Hoang, Pankaj. V. Jadhav, Daniel S. Trettel, Rachel E. Dow, Rachelle M. Baumann, Seokmu Kwon, Kate Matej, Jordan A. Byrne, Christopher A. Azaldegui, Yixuan Liu, Tobias W. Giessen, Hualiang Pi, Shyamal Mosalaganti, Anthony G. Vecchiarelli

## Abstract

Bacteria were once thought to lack organelles, but it is now clear they confine cellular reactions using an array of membrane- and protein-based compartments. A central question, however, is how bacterial organelles are organized in the cell, and whether their spatial control can be engineered. Here, we show that a two-protein system (McdAB) that positions carboxysomes - CO_2_-fixing organelles found in autotrophic bacteria - can be repurposed to provide programmable spatial control to diverse organelles in *Escherichia coli*. McdAB not only restores proper assembly and positioning of heterologously produced carboxysomes in *E. coli*, but can also be reprogrammed to spatially organize all other known types of bacterial organelles, including encapsulins, biomolecular condensates, and even membrane-bound organelles. Programmable spatial organization of bacterial organelles establishes a new design principle for synthetic biology, where the location of reactions is as tunable as their content. Our work paves the way for more efficient biocatalysis in engineered microbes.

---

Cells achieve remarkable control over biochemistry by organizing reactions in space. In bacteria, protein-based microcompartments, encapsulins, and biomolecular condensates, as well as membrane-bound organelles, enable reactions to be concentrated and regulated without the extensive endomembrane systems of eukaryotes. These self-assembling compartments have drawn broad interest as modular building blocks for synthetic biology. Yet, a central challenge limits their utility: when produced in heterologous hosts such as *Escherichia coli*, the nucleoid acts as a diffusion barrier (*1–4*). As a result, without spatial control, these organelles form nucleoid-excluded aggregates at the cell pole. This asymmetric distribution in the cell results in a loss of faithful inheritance across the cell population, compromising activity and constraining their applications.

To solve this problem, most bacteria encode minimal, self-organizing systems comprised of a positioning ATPase from the <u>par</u>tition protein A (ParA) family, along with a partner protein that links the ATPase to its cellular cargo (*5*, *6*). Unlike the linear motors and cytoskeletal systems used by eukaryotes, these two-protein positioning modules generate dynamic ATP-driven protein gradients on the nucleoid to spatially distribute a diversity of cellular cargos, including chromosomes and plasmids (*7*), protein complexes (*8–10*), and organelles (*11*) – ensuring their faithful inheritance and robust functions in the cell. This conserved evolutionary strategy across diverse bacteria and cargo types provides a blueprint for engineering a generalizable method for programming spatial control in bacteria.

The most extensively studied bacterial organelle is the carboxysome - a protein-based organelle found in autotrophic bacteria that is responsible for roughly a third of global CO₂-fixation and holds significant promise for biotechnological applications. Carboxysome organization within the cell is mediated by the two-protein <u>M</u>aintenance of <u>C</u>arboxysome <u>D</u>istribution (McdAB) system (*11*, *12*). McdA is a ParA-family ATPase and McdB is the adaptor protein that links the distribution system to its carboxysome cargo. Specifically, McdB binds carboxysomes and induces McdA oscillations along the nucleoid that distribute carboxysomes throughout the cell to ensure faithful inheritance and CO₂-fixation capacity across the cell population (Supplementary Fig. 1A-B). Without McdAB, carboxysomes mislocalize to the cell poles as nucleoid-excluded aggregates (Supplementary Fig. 1C), leading to asymmetric inheritance and a rapid loss of both carboxysomes and CO₂-fixation capacity in the population (*12–14*).

The McdAB system is currently the only known minimal and self-organizing positioning system for any bacterial organelle. Inspired by its natural anti-aggregation mechanism, we asked whether McdAB could be repurposed to provide synthetic spatial control. Here, we demonstrate that McdAB not only restores proper assembly and positioning of heterologously-produced carboxysomes in *E. coli*, but also generalizes to diverse organelles, including protein-based encapsulins and condensates, and even membrane-bound organelles. Through live-cell imaging, super-resolution microscopy, and *in situ* cryo-electron tomography, we show that McdAB transforms disordered and mispositioned aggregates into uniformly distributed, dynamic arrays. By enabling modular and reversible spatial organization, McdAB provides a versatile strategy for engineering the uniform intracellular distribution of bacterial organelles and assemblies, establishing spatial control as a new design principle for microbial synthetic biology and biotechnology.

## Results

### The McdAB system spatially organizes carboxysomes in E. coli

Carboxysomes are highly efficient CO₂-fixing organelles, and reconstituting them in *E. coli* could transform this industrial microbe into a powerful chassis for sustainable bioproduction. Yet, despite repeated attempts, engineered carboxysomes in *E. coli* remain dysfunctional (*1*, *4*, *15*, *16*). Without their native McdAB positioning system, carboxysomes mislocalize to the cell poles and exhibit poor CO₂-fixation activity. We have previously shown that even in the native autotrophic host, carboxysome mispositioning in the absence of the McdAB system disrupts cell growth and division (*14*). These findings underscore the importance of spatial organization for carboxysome function. Motivated by this, we set out to reconstitute not only carboxysome assembly but also its distribution system in *E. coli*.

We used the recently developed inducible ‘pXpressome’ plasmid toolkit to assemble α- carboxysomes from the chemoautotroph *Halothiobacillus neapolitanus* in *E. coli*. The small subunit of the carbon-fixing Rubisco enzyme (CbbS) was fused to monomeric NeonGreen (mNG) for *in vivo* fluorescence imaging (Fig. 1A; Supplementary Fig. 2A) (*4*). As shown by others, large nucleoid-excluded foci were observed at one or both cell poles (Fig. 1A, C, D; Supplementary Fig. 2A-E). But upon co-expression of the McdAB system, carboxysomes were no longer aggregated at the cell poles (Fig. 1B-D; Supplementary Fig. 2A-E). Instead, multiple foci were distributed across the nucleoid region of the cell. Similar results were obtained in multiple *E. coli* backgrounds, including the more physiologically relevant MG1655 strain and the metabolically engineered *Δrpi* and CCMB1 strains (*17–22*), whose growth depends on Rubisco-mediated carboxylation (Supplementary Fig. 2F). Consistent results with Rubisco fused to either mScarlet or mJuniper (Supplementary Fig. 2G–H) further demonstrate that McdAB-mediated carboxysome positioning is robust to both host genetic background and fluorescent tag.

**Figure 1.**
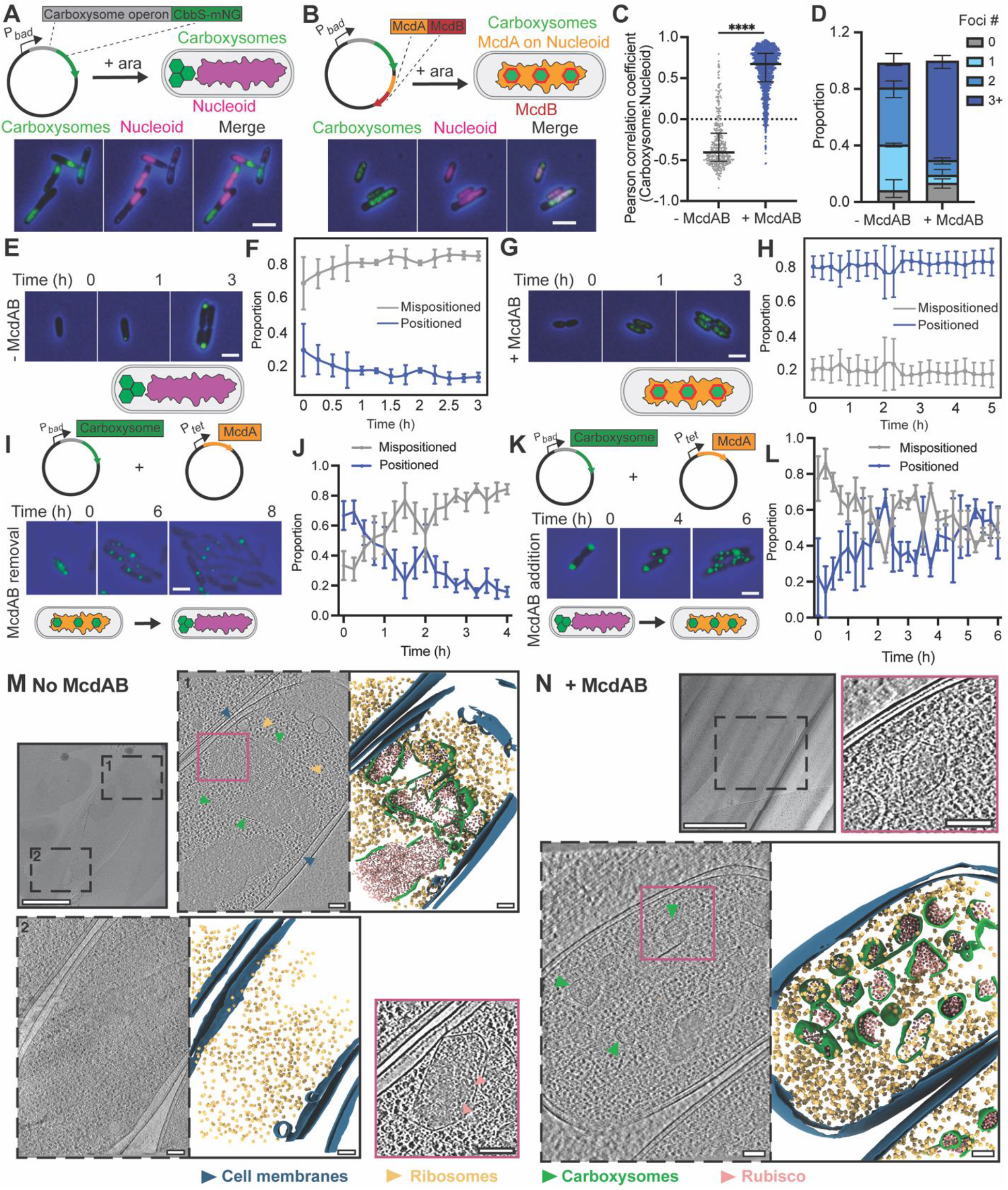
The McdAB system spatially organizes carboxysomes on the *E. coli* nucleoid. (**A**, **B**) Schematic and wide-field fluorescence images of *E. coli* cells assembling carboxysomes without (A) or with (B) the McdAB system. The small subunit of Rubisco (CbbS) is fused to mNeonGreen (mNG); protein expression is induced with 0.2% arabinose (ara). Unless specified, mNG-labeled carboxysomes are shown in green, DAPI-stained nucleoids in magenta, and phase-contrast in blue; scale bar, 2 μm. (**C**) Pearson correlation coefficients measuring colocalization between mNG-carboxysomes and DAPI-stained nucleoids from experiments in A-B. (**D**) Quantification of the proportion of cells with indicated numbers of carboxysome foci from experiments in A-B. (**E**–**H**) Time-lapse analysis showing images and quantification of carboxysome positioning over time without (E, F) or with (G, H) McdAB. Proportions of cells with mispositioned (0–2 polar foci) and positioned (≥ 3 distributed foci) carboxysomes are plotted over time. (**I**–**L**) Reversible control of carboxysome positioning through McdAB removal (I, J) or addition (K, L), as shown by images and quantification of positioned and mispositioned carboxysomes over time. (**M, N**) Cryo-electron tomography of *E. coli* without (M) or with (N) the McdAB system. In each case, low-magnification (6500×) transmission electron microscopy (TEM) images are shown to the left and indicate sites of tomography tilt series data acquisition (dashed boxes). Representative slices through tomograms (dashed boxes) and corresponding segmentations of the tomograms are shown in each case. Cellular features are highlighted as carboxysomes (green), Rubisco (pink), ribosomes (orange), and cell membranes (blue). Zoomed-in views (pink boxes) provide higher-magnification images of carboxysomes (with Rubsico marked with pink arrowheads). The zoomed-in views are shown at different z-heights, where the carboxysome cross-section is maximal. Scale bars: 500 nm (overview), 100 nm (zoom-in).

Having established that McdAB restores carboxysome organization in *E. coli*, we next examined how this positioning system dynamically regulates organelle distribution in growing and dividing *E. coli* cells over time. Cells were classified based on the number and location of foci: “mispositioned” (0–2 polar foci) and “positioned” (≥3 distributed foci). Without McdAB, carboxysome foci remained mispositioned through cell growth and multiple cell divisions (Fig. 1E-F; Video S1). In contrast, approximately 80% of McdAB-expressing cells maintained distributed carboxysome foci (Fig. 1G-H; Video S1). Reducing McdAB levels caused distributed carboxysomes to revert to polar clusters (Fig. 1I-J; Video S2). Conversely, adding McdAB to cells with pre-aggregated carboxysomes triggered their redistribution along the cell length (Fig. 1K-L; Video S2). Expressing McdAB prior to carboxysome production did not enhance their distribution (Supplementary Fig. 2I-J). These results demonstrate that the McdAB system prevents polar aggregation and promotes the even distribution and increased number of α- carboxysome assemblies in *E. coli*.

While fluorescence imaging revealed that McdAB redistributes Rubisco foci across the cell, this approach lacks the necessary spatial resolution to determine whether the observed foci represent correctly assembled carboxysomes. To directly assess the ultrastructure of these assemblies in their native cellular context, we performed *in situ* cryo-electron tomography (cryo-ET) on bacterial cells with and without McdAB. For cells lacking McdAB, tomograms revealed disorganized, densely packed arrays of carboxysome components at the cell poles within the tomograms (Fig. 1M; Supplementary Fig. 3A-E; Video S3). By contrast, and consistent with the fluorescence data, McdAB-expressing cells displayed fully assembled, properly sized, and unclustered carboxysomes distributed across the nucleoid region of the cell volume (Fig. 1N; Supplementary Fig. 3A-E; Video S3). Strikingly, we also observed in our *in situ* cryo-tomograms carboxysome “partitioning,” where the shell appears to invaginate at the midpoint of a larger carboxysome (Supplementary Fig. 3F, white arrowheads), suggesting that McdAB may play a role in carboxysome assembly and size regulation by subdividing oversized structures.

Taken together, these findings provide direct evidence that McdAB is a truly minimal and self-organizing positioning system - requiring only two proteins to distribute carboxysomes in a heterologous host - while simultaneously promoting their structural integrity and proper assembly.

### Spatial control of encapsulin nanocompartments

Building on our discovery that McdAB is minimal and self-organizing, we aimed to repurpose it as a versatile spatial regulator for other bacterial organelles. We recently identified an N-terminal peptide of McdB that is necessary and sufficient for McdA interaction (*23*). Here, we sought to develop this McdB peptide into a <u>m</u>inimal <u>a</u>utonomous <u>p</u>ositioning tag or “MapTag”. By fusing this MapTag to heterologous organelles and co-expressing McdA, we set out to reprogram their distribution in *E. coli*, starting with encapsulins.

While bacterial microcompartments (BMCs), such as the carboxysome, are large and complex structures, encapsulins are much smaller (20-45 nm) and simpler protein-based nanocompartments found in bacteria and archaea (*24*). The shell is composed of one or two self-assembling proteins that tile around an enzymatic cargo loaded via an encapsulin-specific targeting peptide (TP). Like BMCs, encapsulins compartmentalize reaction pathways to sequester toxic intermediates, store reaction products, and increase pathway efficiency, thus offering strategies for specialized metabolic engineering of microbes. No native positioning system has been identified for encapsulins so far.

The study, design, and application of encapsulins rely on their robust production and self-assembly in heterologous hosts like *E. coli*. Previous studies have shown that encapsulins aggregate during assembly in *E. coli* (*2*). Therefore, these encapsulins are ideal substrates for testing the MapTag. Nevertheless, encapsulins have never been visualized in live cells. To address this gap, we first developed a method for live-cell visualization. We engineered an inducible plasmid that expressed an encapsulin shell protein fused to the MapTag, along with a fluorescent cargo protein - mNG fused to a TP (Supplementary Fig. 4A). A degron tag (DT) (*25*) was also fused to mNG to promote degradation of any unencapsulated mNG, thereby enhancing the visualization of mNG- filled encapsulins. Encapsulins without MapTag clustered at the cell poles, consistent with previous studies (*2*) (Supplementary Fig. 4B). Without McdA, the MapTagged encapsulin foci were nucleoid-excluded to one or both cell poles (Fig. 2A, C-D; Supplementary Fig. 2C-D). But when co-expressed with McdA, several encapsulin foci were distributed along the nucleoid (Fig. 2B-D; Supplementary Fig. 4C-D).

**Figure 2.**
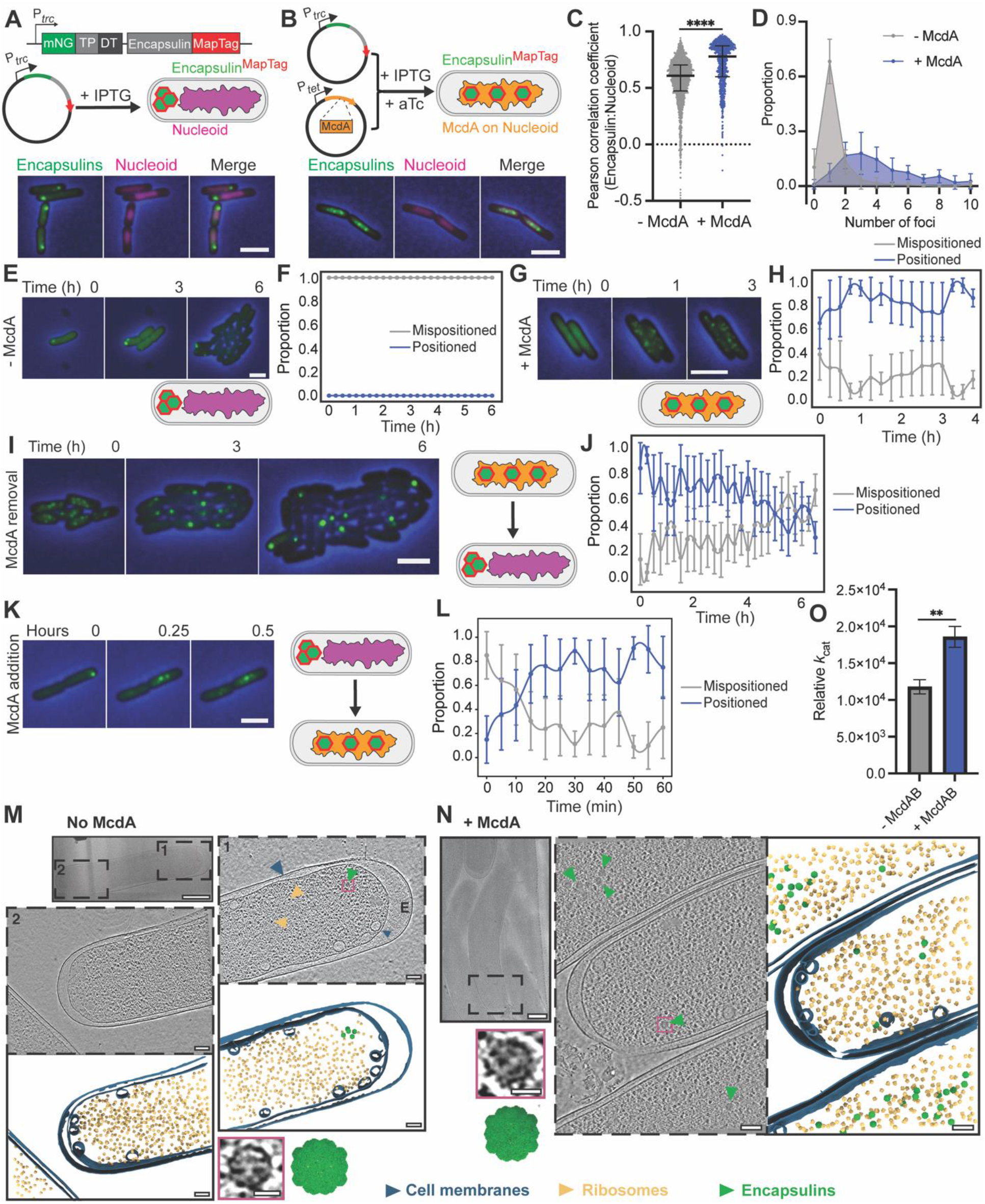
McdA distributes MapTagged encapsulins on the *E. coli* nucleoid. (**A**) Schematic and wide-field fluorescence images of *E. coli* cells assembling encapsulins fused to the MapTag and mNeonGreen (mNG) cargo fused to an encapsulin-specific targeting peptide (TP) and a degron tag (DT), under the P*_trc_*promoter with IPTG induction. (**B**) As in (A), but with McdA co-expressed from a separate plasmid under the P*_tet_* promoter, induced with anhydrotetracycline (aTc). Unless specified, mNG-labeled encapsulins are shown in green, DAPI-stained nucleoids in magenta, and phase-contrast in blue; scale bars, 2 μm. (**C**) Pearson correlation coefficients measuring colocalization between mNG-encapsulins and DAPI-stained nucleoids. (**D**) Quantification of the proportion of cells with the indicated number of encapsulin foci. (**E**–**H**) Time-lapse analysis showing images and quantification of encapsulin positioning over time in cells without (E, F) or with (G, H) McdA; proportions of cells with mispositioned (0–2 polar foci) and positioned (≥3 distributed foci) encapsulins are shown. (**I**–**L**) Reversible control of encapsulin positioning through McdA removal (I, J) or addition (K, L), as shown by images and quantification of positioned and mispositioned encapsulins over time. (**M**, **N**) Cryo-electron tomography of encapsulin^MapTag^-containing *E. coli* without (M) or with (N) McdA. In each case, low-magnification (6500×) transmission electron microscopy (TEM) images are shown to the left and indicate sites of tomography tilt series data acquisition (dashed boxes). Representative slices through tomograms (dashed boxes) and corresponding segmentations of the tomograms are shown in each case. Cellular features are highlighted as encapsulins (green), ribosomes (orange), and cell membranes (blue). Zoomed-in views (pink boxes) provide higher-magnification images of fully assembled encapsulins. Scale bars: 500 nm (overview), 100 nm (zoom-in). (**O**) Relative *k*cat of NanoLuc luciferase in cells containing mispositioned versus positioned encapsulins. NanoLuc, instead of mNG, was targeted to encapsulins, and enzymatic activity was quantified by measuring luminescence following addition of the substrate furimazine.

We next visualized this process in real time. MapTagged encapsulins remained clustered at a single cell pole during the outgrowth of cells lacking McdA (Fig. 2E-F; Video S4). Upon McdA co-expression, encapsulins rapidly redistributed into multiple foci across the cell during outgrowth (Fig. 2G-H; Video S4). Reducing McdA levels caused distributed encapsulins to revert to polar clusters, confirming that positioning requires sustained McdA activity (Fig. 2I-J; Video S5). Conversely, adding McdA to cells with pre-aggregated encapsulins triggered their redistribution along the cell length (Fig. 2K-L; Video S5). Together, these results establish that McdA can spatially control MapTagged encapsulins, with its presence or absence dynamically switching organelles between dispersed and clustered states, respectively.

We further performed cryo-ET analysis to confirm that without McdA, the punctate foci represent fully assembled encapsulin clusters confined to the cell pole (Fig. 2M, Supplementary Fig. 4E, G). By contrast, in McdA-expressing cells, encapsulins were no longer restricted to the poles. Instead, we observed encapsulins distributed along the nucleoid region of the cell (Fig. 2N, Supplementary Fig. 4F-G), consistent with fluorescence imaging.

The results thus far demonstrate that the McdAB system from *H. neapolitanus* can be harnessed for minimal, autonomous spatial control of carboxysomes and encapsulins in *E. coli*. α-carboxysomes are both structurally and phylogenetically distinct from β- carboxysomes found in β-cyanobacteria. To extend the versatility of this system, we developed a complementary β-MapTag derived from the N-terminal region of McdB, a component of the β-carboxysome positioning system of the β-cyanobacterium *Synechococcus elongatus*. Expressing the β-MapTagged-encapsulin genes alongside *S. elongatus* McdA, also resulted in encapsulins being distributed across the *E. coli* nucleoid (Supplementary Fig. 5A-E). Together, these findings establish α- and β- MapTags as broadly applicable systems for programmable spatial control of engineered encapsulin nanocompartments.

To determine whether spatial control influences the activity of an encapsulated enzyme, we replaced mNG with NanoLuc luciferase as the encapsulin cargo. In the presence of McdA, both α- and β-MapTagged encapsulins exhibited nearly twofold increases in relative *k*cat and endpoint NanoLuc activity compared with cells lacking McdA (Fig. 2O, Supplementary Fig. 5F-G). Thus, MapTag-mediated spatial organization not only controls encapsulin distribution but also enhances encapsulated enzyme activity, demonstrating that organelle positioning can be harnessed to improve the performance of engineered nanocompartments in bacteria.

### Spatial control of biomolecular condensates

While BMCs and encapsulins have long been recognized as bacterial organelles, biomolecular condensates have only recently emerged as a distinct strategy for organizing cellular reactions in bacteria (*26*). Their dynamic and reversible nature makes condensates attractive for biotechnology (*27*), yet in bacteria, they typically coalesce into a single polar focus, leading to asymmetric inheritance during division (*3*, *28*). To address this limitation, we engineered a condensate positioning system using the MapTag.

*S. elongatus* McdB forms nucleoid-excluded condensates when expressed in *E. coli* (*28*). We first asked whether McdA could redistribute these condensates on the nucleoid. When expressed alone, mNG-McdA colocalized with the *E. coli* nucleoid (Supplementary Fig. 6A-B). Intriguingly, co-expression with McdB abolished this association. Because McdB condensates are highly fluid-like and a substantial fraction remains cytoplasmic (*28*), we hypothesized that this cytoplasmic McdB pool globally stimulates McdA release from the nucleoid. To reduce spurious interactions away from the condensate cargo, we fused McdB to the PopTag, a well-characterized synthetic biomolecular condensate scaffold derived from the *Caulobacter crescentus* polar organizing protein PopZ (*29*). PopTag has been widely used as a modular heterologous condensate scaffold in bacteria (*28*), making it an ideal system for testing whether the MapTag–McdA module can spatially organize biomolecular condensates unrelated to the native Mcd positioning system. Fusion of McdB to PopTag (McdB^PopTag^) enhanced McdB sequestration into condensates (Supplementary Fig. 6C-D). Without McdA, McdB^PopTag^ formed one or two large polar condensates (Fig. 3A, C-D; Supplementary Fig. 7A). But upon McdA co-expression, multiple condensates were distributed along the nucleoid (Fig. 3B, C-D; Supplementary Fig. 7A).

**Figure 3.**
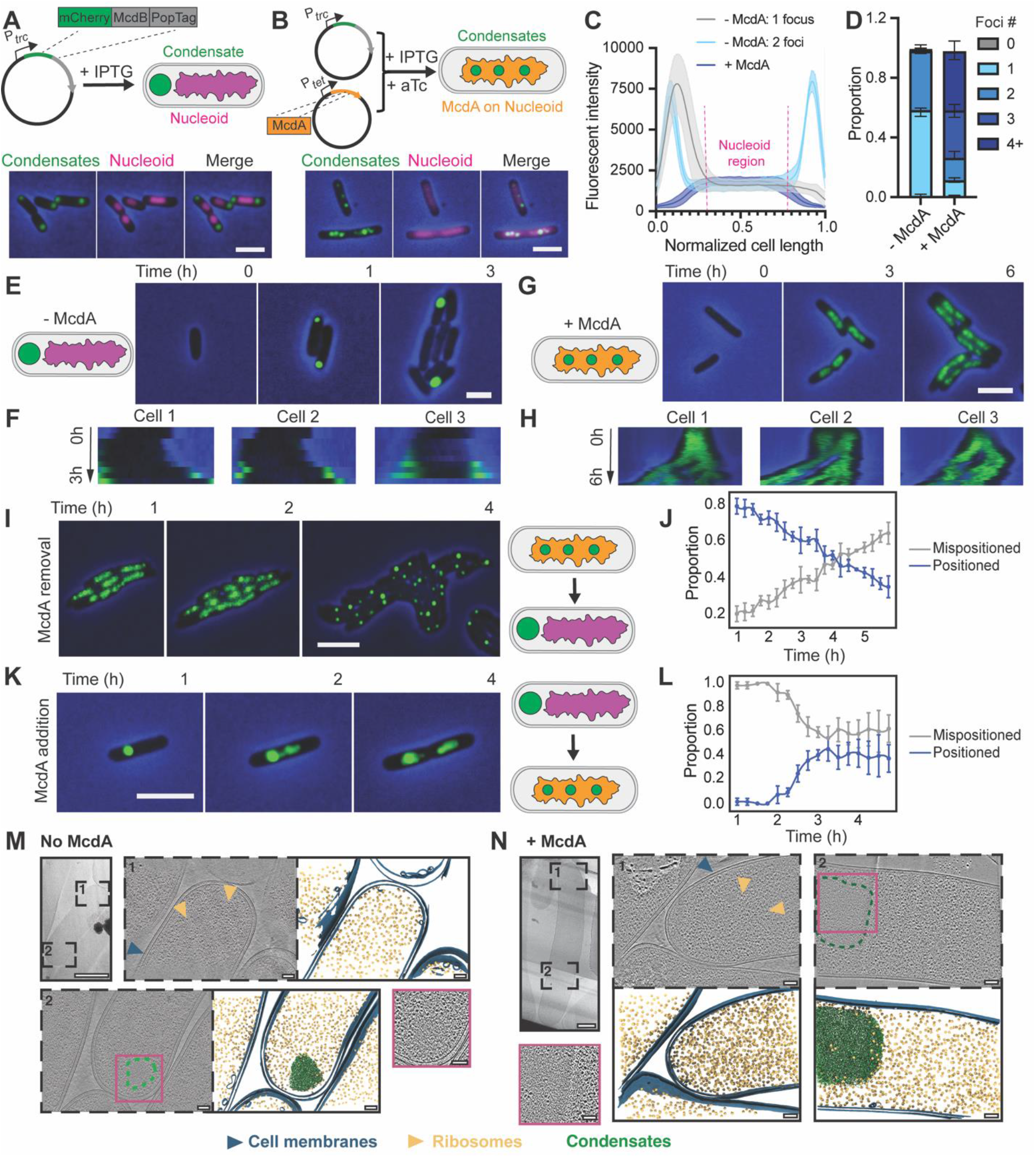
McdA distributes McdB condensates on the *E. coli* nucleoid. (**A**) Schematic and wide-field fluorescence images of *E. coli* cells expressing McdB condensates fused to mCherry and PopTag, under the P*_trc_* promoter with IPTG induction. (**B**) As in (A), but with McdA co-expressed from a separate plasmid under the P*_tet_* promoter, induced with anhydrotetracycline (aTc). Unless specified, McdB^PopTag^ condensates are shown in green, DAPI-stained nucleoids in magenta, and phase-contrast in blue; scale bars, 2 μm. (**C**) Fluorescent intensity profiles of McdB^PopTag^ signal along the normalized cell length. (**D**) Quantification of the proportion of cells with the indicated number of McdB^PopTag^ condensates from experiments in A-B. (**E–H**) Time-lapse analysis showing wide-field fluorescence images (E, G) and kymographs of representative cells (F, H) depicting the distribution of McdB^PopTag^ fluorescence over time without (E, F) or with (G, H) of McdA. (**I–L**) Reversible control of McdB^PopTag^ condensate positioning by McdA removal (I, J) or addition (K, L), shown by wide-field fluorescence images and quantification of mispositioned (0–2 polar foci) and positioned (≥3 distributed foci) condensates over time. (**M**, **N**) Cryo-electron tomography of *E. coli* assembling McdB^PopTag^ condensates without (M) or with (N) McdA. In each case, low-magnification (6500×) transmission electron microscopy (TEM) images are shown to the left and indicate sites of tomography tilt series data acquisition (dashed boxes). Representative slices through tomograms (dashed boxes) and corresponding segmentations of the tomograms are shown in each case. Cellular features are highlighted as condensates (green), ribosomes (orange), and cell membranes (blue). Zoomed-in views (pink boxes) provide higher-magnification images of condensates seen as ribosome-excluded areas. Scale bars: 500 nm (overview), 100 nm (zoom-in).

Without McdA-dependent positioning, polar condensates were rapidly lost from the cell population (Fig. 3E-F; Video S6). By contrast, in the presence of McdA, all daughter cells inherited roughly an equal complement of McdB^PopTag^ condensates distributed over the nucleoid (Fig. 3G-H; Video S6). Reducing McdA levels over time resulted in distributed condensates that coalesced into a single, large nucleoid-excluded condensate at the cell pole (Fig. 3I-J; Video S7). Conversely, we observed that replenishing McdA led to the dissolution of polar condensates and the formation of multiple condensates distributed along the nucleoid (Fig. 3K-L; Video S7). Finally, *in situ* cryo-ET confirmed the relocalization of polar condensates to the nucleoid in the presence of McdA by *in situ* cryo-ET (Fig. 3M-N, Supplementary Fig. 7B-D). Importantly, our cryo-ET analysis revealed that the addition of tags and the positioning system did not alter the property of the PopTag to form condensates.

After successfully engineering a system that allowed us to distribute full-length β-McdB condensates, we set out to identify the minimal β-MapTag sequence for spatially regulating heterologous condensates with McdA. The first 20 N-terminal amino acids of β-McdB fused to mCherry^PopTag^ did not yield any condensates (Supplementary Fig. 8A-B). However, addition of the adjacent coiled-coil dimerization domain of β-McdB provided additional valency that restored condensate assembly (Supplementary Fig. 8C). Co-expression of McdA distributed these β-MapTagged condensates along the nucleoid (Supplementary Fig. 8D-F). When swapping the β-MapTag for the α-MapTag, co-expression with its cognate McdA resulted in condensate dissolution over the nucleoid (Supplementary Fig. 8G-I), suggesting the α-MapTag may be more suitable for positioning condensates with lower saturation concentrations. Together, we demonstrate that McdA can direct the spatial organization of both McdB condensates and MapTagged heterologous condensates, ensuring their cellular distribution and faithful inheritance across the cell population.

### Spatial control of a membrane-bound organelle

To test the versatility and robustness of MapTags, we determined if this mode of spatial control could be extended beyond protein-based compartments to include membrane-bound organelles. Ferrosomes are membrane-bound compartments for iron storage that require the membrane protein FezA and the P1B6-ATPase transporter FezB for their formation (*30*). Heterologous expression of mNG-FezB fused to the α-MapTag, along with untagged FezA in *E. coli*, resulted in the formation of ferrosome clusters at the cell pole (Fig. 4A, C-D; Supplementary Fig. 9A-B). Upon co-expression of McdA, multiple ferrosome foci were observed over the nucleoid (Fig. 4B, C-D; Supplementary Fig. 9A-B). A co-expression time course showed that McdA facilitates the cellular distribution and faithful inheritance of assembling ferrosomes across the cell population (Fig. 4E-F). We observed that McdA was also able to partition preformed ferrosome clusters at the cell pole and redistribute them over the nucleoid (Fig. 4G-H). Finally, we observed that fusing the β-MapTag to FezB was just as effective at distributing ferrosomes as the α-MapTag (Supplementary Fig. 9C-D).

**Figure 4.**
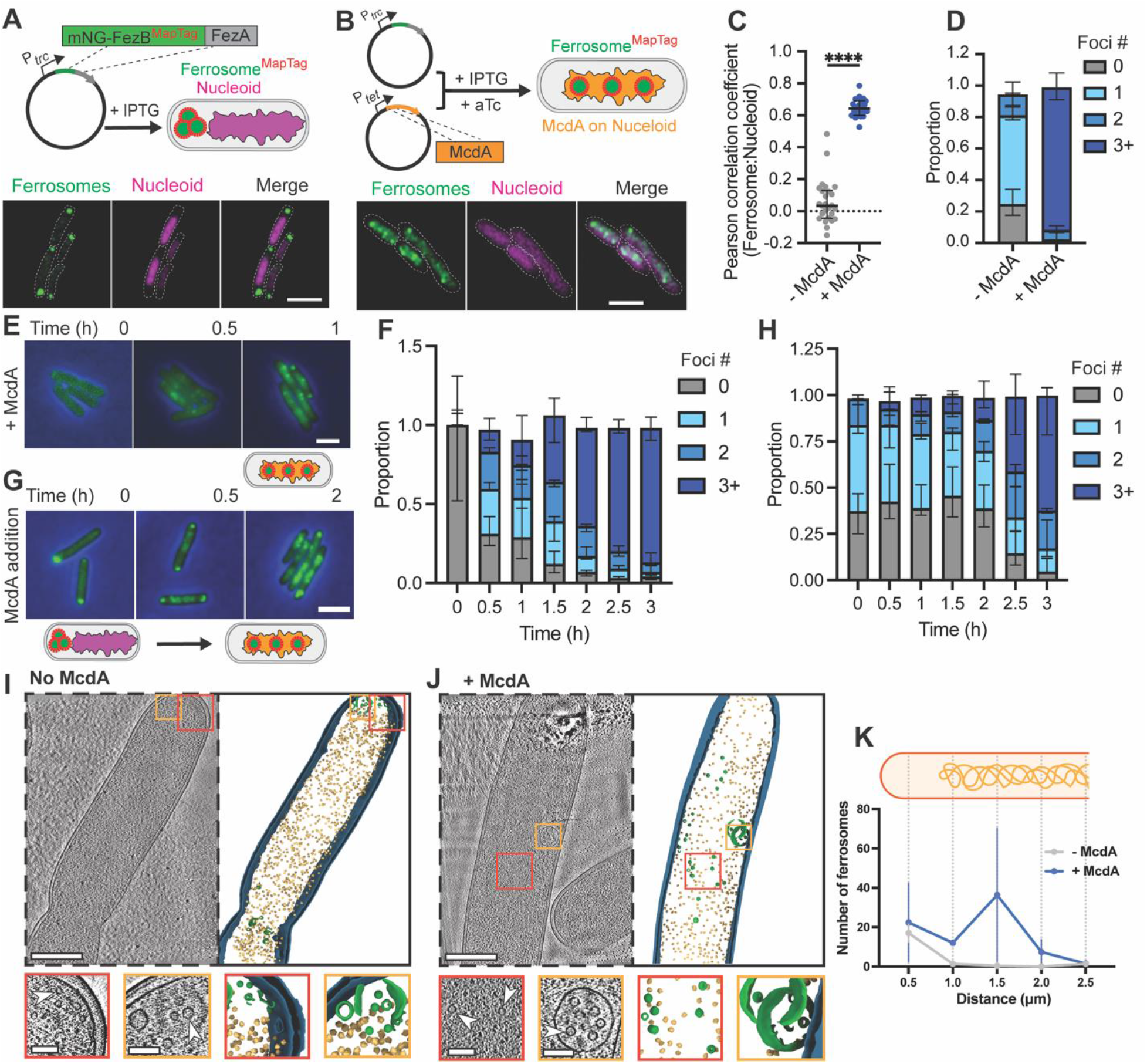
McdA distributes MapTagged ferrosomes on the *E. coli* nucleoid. (**A**) Schematic and super-resolution Airyscan images of *E. coli* cells assembling ferrosomes, composed of FezA and FezB (with FezB fused to mNeonGreen [mNG] and the MapTag), under the P*_trc_* promoter with IPTG induction. (**B**) As in (A), but with McdA co-expressed from a separate plasmid under the P*_tet_* promoter, induced with anhydrotetracycline (aTc). Ferrosomes are shown in green and DAPI-stained nucleoids in magenta,. Scale bars, 2 μm. (**C**) Pearson correlation coefficients measuring colocalization between mNG-labeled ferrosomes and DAPI-stained nucleoids. (**D**) Quantification of the proportion of cells with the indicated number of ferrosome foci. (**E**, **F**) Wide-field fluorescence images (E) and quantification (F) of ferrosome positioning in E over a 3-h time course in cells with McdA. Ferrosomes are shown in green and phase contrast in blue. (**G**, **H**) Wide-field fluorescence images (G) and quantification (H) showing the redistribution of ferrosomes in G upon McdA addition to cells initially lacking McdA over a 3-h time course. (**I**-**J**). Cryo-electron tomography of ferrosome^MapTag^-containing *E. coli* cells without (**I**) or with (**J**) McdA. Representative slices through tomograms (dashed boxes) and corresponding segmentations are shown for each condition. Cellular features are highlighted as ferrosomes (green), ribosomes (gold), and cell membranes (teal). Zoomed-in views (red and orange boxes) show higher-magnification images of representative ferrosomes (arrowheads) in both raw tomogram and segmented reconstructions. Scale bars: 500 nm (overview), 100 nm (zoom-in). (**K**) Quantification of ferrosome number as a function of distance from the cell pole. The schematic above the graph illustrates the positional bins used for quantification along the cell long axis.

We further performed cryo-ET analysis to confirm that without McdA, the punctate foci represent ferrosome clusters confined to the cell pole (Fig. 4I, K). By contrast, in McdA- expressing cells, ferrosomes were no longer restricted to the poles. Instead, ferrosomes distributed along the nucleoid region of the cell (Fig. 4J-K), consistent with fluorescence imaging.

All fluorescent cargo probes used in this study were expressed predominantly as full-length proteins with minimal degradation (Supplementary Fig. 10), indicating that the observed localization patterns faithfully represent the intracellular distribution of the cargo.

## Discussion

We establish uniform spatial organization as a new design principle in bacterial synthetic biology. By reconstituting and repurposing McdAB, we show that a minimal, two-protein positioning module can be harnessed to restore the native organization of carboxysomes and, more broadly, to spatially reprogram the distribution of diverse bacterial organelles in *E. coli*. The capacity to engineer both the composition of cellular compartments and their uniform intracellular distribution establishes spatial organization as a programmable design parameter in bacterial synthetic biology.

The impact of McdAB-mediated spatial control extends beyond simple organelle placement. We demonstrate that mislocalized carboxysomes in *E. coli* also suffer structural defects. McdAB corrects this by transforming disordered polar aggregates into properly sized and structurally intact organelles distributed over the nucleoid, restoring both assembly fidelity and inheritance. By generalizing this approach with MapTags, we extend spatial control to encapsulins, condensates, and ferrosomes, spanning the full spectrum of known protein-based and lipid-bound organelle types. These results suggest that spatial misregulation may be a universal bottleneck for heterologous organelle engineering, and that positioning systems provide a general solution.

This conceptual advance parallels the way cytoskeletal elements and trafficking systems underpin compartmentalization in eukaryotes. Whereas eukaryotes evolved complex, multi-component transport machinery, bacteria have distilled spatial organization into minimal ATP-driven modules that establish self-organizing gradients. By reengineering this strategy, we unlock a scalable, orthogonal method for positioning organelles in bacterial cells. The versatility of α- and β-MapTags across structurally distinct compartments underscores their potential as modular “address labels” for intracellular engineering.

The α- and β-MapTag systems presented here constitute a small subset of bacterial organelle positioning modules. Many bacterial organelles and protein complexes are spatially organized by distinct ParA-family ATPases and their cognate adaptor proteins, raising the possibility that additional orthogonal positioning systems can be identified and repurposed. Expanding this repertoire could enable the simultaneous spatial organization of multiple engineered organelles within the same cell, allowing independent positioning of distinct metabolic pathways or reaction centers while minimizing crosstalk between positioning circuits. Such multiplexed spatial programming would establish a new level of intracellular organization for engineered bacteria.

Programmable organelle positioning opens several future directions. First, it enables the rational design of metabolic “reaction landscapes” in bacteria, where organelles can be spaced to prevent interference, aligned to channel substrates, or dynamically repositioned in response to environmental signals. Second, because McdAB activity is reversible, spatial organization can be toggled on or off, allowing for time-dependent control of metabolism or stress responses. Third, distributing compartments evenly across the nucleoid ensures faithful inheritance, reducing population-level heterogeneity and thereby improving the reliability of engineered microbial systems. By layering spatial control onto existing advances in compartment design, we foresee the emergence of “organelle engineering” as a discipline in its own right - where the content, function, and location of organelles can all be rationally specified. Such a framework could accelerate the development of microbial cell factories for carbon capture, green chemistry, and therapeutics.

Beyond biotechnology, this work provides a blueprint for constructing synthetic cells. A central question in bottom-up biology is how to couple self-assembly with inheritance. Our results demonstrate that a minimal gradient-based system is sufficient to transform self-assembling but static compartments into dynamic, heritable organelles.

## Acknowledgements

We thank Dr. Eric Rentchler and Dr. Sasha Meshinchi of the Biomedical Microscopy Core at the University of Michigan for training and coordinating access to the super-resolution Airyscan Microscope. A patent application related to this work (no. 19/527,740) has been filed.

## Funding

National Science Foundation grant 1941966 (A.G.V)

National Institutes of Health grant R35GM152128 (A.G.V.)

National Institutes of Health 1K99GM157383 and 4R00GM157383-03 (Y.H.)

National Institutes of Health 1DP2GM150019-01 (S.M.)

Klatskin Sutker Discovery Fund (S.M.)

National Institutes of Health grant S10OD030275 (S.M.)

Arnold and Mabel Beckman Foundation award to the University of Michigan Cryo-EM facility

National Institutes of Health grant R35GM133325 (T.W.G.)

National Science Foundation grant 2342136 (T.W.G.)

National Institutes of Health DP2AI184552 (H.P.)

Searle Scholar Award SSP-2025-109 (H.P.)

National Institutes of Health T32 GM145304 (J.A.B.)

National Science Foundation GRFP (R.M.B.)

University of Michigan Rackham Graduate Student Research Grant (J.A.B. and R.M.B.)

Rackham Predoctoral Fellowship and Asan Foundation Biomedical Science Scholarship (S.K.)

The funders had no role in the study design, data collection, analysis, or the content and publication of this manuscript.

## Author contributions

Conceptualization: A.G.V. and Y.H.

Methodology: A.G.V., Y.H., P.V.J., D.S.T., R.E.D., R.M.B., S.K., K.T., Y.L., J.A.B., C.A.A., T.W.G., H.P., and S.M.

Investigation: A.G.V., Y.H., P.V.J., S.M.

Visualization: A.G.V., Y.H., R.E.D., J.A.B.

Funding acquisition: A.G.V., Y.H., S.M., T.W.S., H.P.

Project administration: A.G.V. and Y.H.

Supervision: A.G.V. and S.M.

Writing – original draft: A.G.V. and Y.H.

Writing – review & editing: All authors

## Data and materials availability

All plasmids are available at Addgene. Data processing and analysis scripts for this study are available on GitHub.

## Supplementary Information

## Materials and Methods

### Plasmid construction

Primers, plasmids, and strains used in this study are listed in Supplementary Tables 1 & 2. Plasmids were cloned using Gibson assembly and transformed into competent *E. coli* DH5ɑ cells, and transformants were selected on LB plates supplemented with the corresponding antibiotics.

#### Carboxysomes

Plasmids encoding fluorescently tagged CbbS with McdAB were generated using Gibson assembly. For example, to create pDT081, the previously published carboxysome-only construct pDT074 (*1*) was PCR-linearized with Q5 polymerase and primers oDT077/oDT098 (15 s/kb extension, 65°C annealing, 27 cycles). The 13.7 kb amplicon was treated with DpnI (37°C, 30 min), purified (Zymo PCR Clean and Concentrator Kit), and eluted in deionized water. The *H. neapolitanus* McdAB cassette (gDT062; Twist Biosciences), codon-optimized for *E. coli* and containing each gene preceded by the strong BBa_0030 ribosome binding site, was assembled with 0.020 pmol linearized vector and 0.060 pmol gDT062 using the NEBuilder HiFi DNA Assembly kit (50°C, 20 min). Gibson reactions were immediately frozen at −20°C and transformed into NEB5alpha competent cells by heat shock. Colony screening was performed by whole-plasmid sequencing (Plasmidsaurus). Plasmids pDT082 and pDT084 were constructed from the published plasmids pDT075 and pDT078 using a similar strategy.

#### Encapsulins

Plasmid pSMK1, for the expression of encapsulin fused to mNeonGreen and an α- MapTag, was generated by Gibson assembly of a pTrc99A backbone (*2*) with a synthetic, *E. coli* codon-optimized gene fragment encoding mNeonGreen and α-McdB-tagged encapsulin. Plasmid pSMK2 was prepared by replacing the α-McdB tag in pSMK1 with a β-MapTag, using overhang PCR with primers pSMK23-F and pSMK2-R. Plasmid pYH84, used to express McdA from *H. neapolitanus*, was constructed by Gibson assembly of a pBbA2k backbone (*3*) with a synthetic, *E. coli* codon-optimized gene fragment encoding *H. neapolitanus* McdA. Plasmid pYH91, for expression of McdA from *S. elongatus*, was generated using the same strategy with a corresponding gene fragment.

#### Condensates

Plasmid pYH82, which expresses the McdB^PopTag^ condensate, was constructed by Gibson assembly. The backbone was amplified from pYH77 (*2*) using primers YH20 and YH21, and the *S. elongatus* McdB fragment was amplified from pYH71 using primers YH22 and YH23. The plasmid pJAB30 was constructed by amplifying pYH82 with primers YH24 and YH25, followed by ligation with KLD Enzyme Mix (New England Biolabs), transformation into *E. coli* DH5α, and plasmid verification by Plasmidsaurus. The plasmid pJAB39 was generated using a similar strategy with primers YH26 and YH27.

#### Ferrosomes

To construct pKM1, pSMK1 was sequentially digested with NcoI-HF and SalI-HF at 37°C for one hour per enzyme, and the plasmid backbone was purified using a gel extraction kit. A GeneBlock containing α-MapTag-mNeonGreen was obtained from Integrated DNA Technologies and amplified with primers KM1/KM2. FezA and FezB were amplified from *C. difficile* CD196 genomic DNA using the primers KM3/KM4 and KM5/KM6, respectively. PCR products were purified (Qiagen PCR purification kit) and assembled with the backbone in a four-piece HiFi DNA Assembly reaction, then transformed into *E. coli* DH5α. Resulting transformant were verified by whole-plasmid sequencing (Plasmidsaurus). For construction of pKM2, a GeneFragment encoding the β-MapTag sequence was obtained from Genewiz. pKM1 and the GeneFragment were digested with KasI and BamHI-HF at 37°C for one hour, then purified via gel extraction. The fragments were ligated with T4 DNA ligase at room temperature for 16 hours and transformed into *E. coli* DH5α. Plasmids were mini-prepped and correct insertion was verified by Sanger sequencing (Keck DNA Sequencing Facility, Yale University).

### Media and culture growth

For routine bacterial growth, LB medium was used for all overnight cultures. Single colonies were inoculated into 15 mL Falcon tubes containing LB broth and grown overnight at 37°C with agitation (200-225 rpm) for 12-16 hours. Exponential phase cultures were initiated by diluting overnight cultures 1:100 into LB or AB minimal medium. Carboxysome production experiments in *E. coli* Top10 were performed in LB medium. Experiments in *E. coli* MG1655 were carried out in AB minimal medium. AB medium was supplemented with 0.2% carbon source (glycerol for growth or glucose to minimize basal expression from the P*_trc_* promoter), 0.2% casamino acids, 10 μg/mL thiamine, and 25 μg/mL uracil (*4*). Where appropriate, antibiotics were added at the following final concentrations: carbenicillin (100 μg/mL), kanamycin (100 μg/mL), and chloramphenicol (100 μg/mL). Induction of protein production was achieved using IPTG, anhydrotetracycline (aTc), or arabinose at concentrations detailed in each experiment. Strains and plasmids are listed in Table S1. Reagents are listed in Supplementary Table 3. All media compositions and induction times are detailed in Supplementary Table 3.

### Expression of organelles

#### Carboxysomes

*E. coli* Top10 cells were transformed with plasmid pDT074 to express mNeonGreen-labeled carboxysomes. For the respective carboxysomes with the McdAB system, pDT81 was used. Overnight cultures were diluted and grown in LB medium until reaching an optical density (OD₆₀₀) of 0.2–0.6. Carboxysome expression was induced with 0.2% arabinose for 2 hours. For fluorescently labeled variants, pDT075 and pDT082 were used for mScarlet-labeled carboxysomes, and pDT078 and pDT084 for mJuniper-labeled carboxysomes.

#### Encapsulins

*E. coli* MG1655 *ΔlacY* cells were transformed with plasmid pSMK1 to express encapsulin^α-MapTag^. Exponential phase cultures in AB media were induced with 1 mM IPTG for 2 hours. For co-expression of McdA, cells were co-transformed with pSMK1 and pYH84 and induced with 1 mM IPTG and 20 nM aTc. For encapsulin^β-MapTag^, plasmids pSMK2 and pYH91 were used under similar induction conditions.

#### Biomolecular condensates

To generate a heterologous synthetic condensate, we fused *S. elongatus* McdB to PopTag, a synthetic condensate scaffold derived from the *Caulobacter crescentus* polar organizing protein PopZ (*5*). PopTag promotes robust condensate formation in *E. coli*, allowing us to test whether the MapTag–McdA positioning module can spatially organize biomolecular condensates independently of the native Mcd system. For full-length McdB^PopTag^ condensates, *E. coli* MG1655 *ΔlacY* cells were transformed with plasmid pYH82. Exponential phase cultures in AB medium were induced with 100 μM IPTG for 2 hours. For co-expression with McdA, cells were co-transformed with pYH82 and pYH91 and induced with 100 μM IPTG and 5 nM aTc. For α- and β-MapTag-tagged condensates, cells were co-transformed with pJAB39 and pYH82, or with pRED2 and pYH84, respectively, and induced under the same conditions.

#### Ferrosomes

For ferrosome^α-MapTag^, *E. coli* MG1655 *ΔlacY* cells were transformed with plasmid pKM1. Exponential phase cultures in AB medium were induced with 1 mM IPTG for 2 hours. For co-expression with McdA, cells were co-transformed with pKM1 and pYH84 and induced with 1 mM IPTG and 20 nM aTc. For ferrosome^β-MapTag^, cells were co-transformed with pKM2 and pYH91 and induced under the same conditions.

#### McdA removal and addition assays

To reduce McdA protein levels in cells with positioned organelles, cultures were washed (5,000 rpm for 2 minutes, three times) and resuspended in fresh medium containing IPTG but lacking aTc, the inducer for all McdA expressing plasmids. As cells grew and divided, McdA levels decreased by generational dilution. Cells were monitored by live-cell fluorescent imaging. In a complementary experiment, cells expressing both MapTagged organelle and McdA plasmids were initially induced with IPTG (1 mM) in the absence of aTc, resulting in aggregated, mispositioned compartments. Upon addition of aTc (20 nM) and subsequent expression of McdA to these cultures, time-lapse fluorecent imaging was performed to monitor compartment redistribution and positioning.

#### DAPI staining and chloramphenicol treatment

DAPI was added to exponentially growing cultures at a final concentration of 2 μM and incubated for 15 min at 37°C before imaging, without rinsing. DAPI fluorescence was detected using the “DAPI” filter set as described in the “Wide-field fluorescence and phase-contrast imaging” section.

Because the nucleoid occupies most of the cell volume, it is difficult to assess the extent of McdA–nucleoid colocalization. Nucleoid compaction facilitates the determination of whether McdA is associated with the nucleoid or freely diffuses throughout the cytoplasm. For nucleoid compaction, chloramphenicol was added to exponentially growing cultures at a final concentration of 100 μg/mL, followed by incubation at 37°C for 30 min before imaging.

#### NanoLuc enzymatic assay in whole cell

NanoLuc activity was quantified by measuring luminescence following addition of its substrate, fluorofurimazine, using a Synergy H1 plate reader. *E. coli* cells producing the indicated constructs (α-MapTag or β-MapTag, with or without McdA) were induced with IPTG (1mM) for 2 hours and diluted in minimal media to a final OD₆₀₀ of 0.1. Fluorofurimazine substrate was added at a single final concentration of 10 μM. Cell suspensions were dispensed into flat white bottom nonbinding surface 96-well plates (Corning 3990) in a final reaction volume of 100 μL, with AB minimal media alone serving as a background control. All conditions were assayed in triplicate at room temperature. Luminescence was monitored at 1-minute intervals following substrate addition, and background luminescence (minimal media without cells) was subtracted from each sample reading.

Relative *k*cat was determined by linear regression of the background-subtracted luminescence signal over the first 0–10 minutes following substrate addition, with the resulting slope reported as relative *k*cat. Endpoint luminescence was recorded as the background-subtracted signal averaged over the 20–30 minute time window of the assay.

#### Wide-field fluorescence and phase-contrast imaging

Wide-field fluorescence and phase-contrast imaging were performed using a Nikon Ti2-E motorized inverted microscope controlled by the NIS Elements software with a SOLA 365 LED light source, a 100 × objective lens (Oil CFI Plan Apochromat DM Lambda Series for Phase Contrast), and a Photometrics Prime 95B back-illuminated sCMOS camera or a Hamamatsu Orca-Flash 4.0 LTS camera. Fusions to mNeonGreen and mJuniper were imaged using a “GFP” filter set (C-FL GFP, Hard Coat, High Signal-to-Noise, Zero Shift, Excitation: 436 ± 10 nm, Emission: 480 ± 20 nm, Dichroic Mirror: 455 nm). DAPI fluorescence was imaged using a “DAPI” filter set (C-FL DAPI, Hard Coat, High Signal-to-Noise, Zero Shift, Excitation: 350 ± 25 nm, Emission: 460 ± 25 nm, Dichroic Mirror: 400 nm). mScarlet and mCherry were imaged using a “Texas Red” filter set (C-FL Texas Red, Hard Coat, High Signal-to-Noise, Zero Shift, Excitation: 560 ± 20 nm, Emission: 630 ± 37.5 nm; Dichroic Mirror: 585 nm).

#### Time-lapse videos of the induction of protein production

To visualize the dynamics of organelle positioning, exponential-phase cultures were induced with either IPTG alone or with both IPTG and aTc. After incubation at 37 °C with shaking at 200 rpm for 15 min, a 2 μL aliquot of culture was then spotted onto agarose pads (LB medium for carboxysomes; AB medium for encapsulins and condensates) in 35-mm glass-bottom dishes, with cells positioned between the agar pad and the glass surface. Time-lapse images were captured using the phase contrast and “GFP” filter set (see "Wide-field fluorescence and phase-contrast imaging" section) every 15 min for 3 hours. For McdA removal experiments, cultures were washed (5,000 rmp for 2 min, three times), resuspended in fresh medium lacking aTc, and spotted onto AB agarose pads without aTc. For McdA addition experiments, cultures 2 hours post-IPTG induction were spotted onto AB agarose pads containing 20 nM aTc.

#### Super-resolution imaging

Cells were spotted onto agar pads in glass-bottom dishes for imaging. Super-resolution images were acquired using a Zeiss LSM 980 Airyscan 2 microscope equipped with a 63×/1.4 NA Plan Apo oil objective (190 μm working distance). mNeonGreen fluorescence was excited with a 488 nm laser and detected with the Airyscan 2 detector. Images were post-processed using Zen 3.4 (Blue edition).

#### Cryo-electron tomography (cryo-ET) of *E. coli*

##### Sample vitrification

Bacterial samples were first concentrated to an OD_600_ of ∼40 by centrifugation at 500 × g for 5 min. Subsequently, 3 μL of the concentrated bacterial cell suspension was applied to glow-discharged R1/4, 200 mesh holey copper grids (Quantifoil Micro Tools GmbH). Grids were single-side blotted for 5 s with a blot force of 10 at 30 °C and 95% humidity before being plunge-frozen in liquid ethane using Vitrobot Mark IV (ThermoFisher Scientific), clipped onto AutoGrids (ThermoFisher Scientific), and stored in liquid N_2_ till further use.

#### Cryo-focused ion beam milling (cryo-FIB) and tilt series acquisition

Clipped grids were loaded onto an Aquilos 2 dual-beam microscope (Thermo Fisher Scientific), maintained at ≤−190 °C. Grids were coated with a layer of organometallic platinum using a gas injection system (GIS) for 30 s and subsequently sputter-coated for 20s. The lamella milling sites were selected using the MAPS 3.25 software (Thermo Fisher Scientific). At each chosen site, tension-relief trenches were milled with a beam current of 1 nA. Subsequently, ∼5 µm-thick lamellae were milled at an angle of 6 degrees. Following this, lamellae were thinned stepwise to ∼1 µm thickness with beam currents of 500–100 pA. Finally, all lamellae were polished to a final thickness of ∼200 nm with a beam current of 50 pA. Tilt series were acquired on a Titan Krios G4i transmission electron microscope (Thermo Fisher Scientific) operated at 300 kV, equipped with a K3 direct-electron detector (Gatan Inc.) and a Bioquantum energy filter (Gatan Inc.). Montages of lamella were collected at a magnification of 6500× to identify cells with features of interest. Movies were collected at two different magnifications of 33,000x (high magnification) and 19,500x (low magnification) in super-resolution mode and binned 2×, corresponding to a nominal physical pixel size of 2.654 Å and 4.504 Å, with 9 frames per tilt. Tilts series were collected using SerialEM software v4.1 (*6*) with a dose-symmetric tilt scheme (*7*) with 3° tilt increment from -60 ° to +60°, starting from a + 6° pretilt. The total dose per tilt series was 120 e−/Å^2^.

### Tilt series pre-processing, alignment, and tomogram reconstruction

Motion correction and contrast transfer function (CTF) estimation were performed using WarpTools v2.0.0 . Subsequently, motion-corrected tilts were aligned using patch tracking in batch mode using a custom bash script and, if required, optimized manually in IMOD v4.11.25 (*6*) . The resulting alignment files were imported into Warp. 4× binned tomograms, resulting in a pixel size of 10.60 Å (high magnification) and 18.02 Å (low magnification), were reconstructed using WarpTools and used for downstream analysis.

### Membrane tracing and voxel segmentation

The binned tomograms were processed with Membrain-Seg (*8*) for tracing and segmenting cellular membranes and carboxysomes. The traced volumes were imported into AMIRA (Thermo Fisher Scientific) for manual curation. Next, smooth surfaces were generated from the membrane and carboxysome labels. These surfaces were imported into ChimeraX (*9*) for final rendering. Template matching was performed using WarpTools to segment ribosomes using the *E. coli* 70S ribosome density map (EMD-13270) (*10*) as the template. The resulting particle picks were manually curated to get the final ribosome particle coordinates. These coordinates were used to place ribosomes in ChimeraX using the ArtiaX v0.6.0 plugin (*11*) . Rubisco and encapsulins were manually picked in the tomograms, and the particle coordinates were imported into ChimeraX to place the Rubisco (EMD-15409) (*12*) and encapsulin (EMD-5917) (*13*) density maps into the volumes.

### Quantification of organelle distribution

Low-magnification tomograms were used to quantify organelle distributions in both positioned and non-positioned samples. For carboxysomes, encapsulins and ferrosomes, organelle distribution was calculated by counting the number of organelles per 0.5 μm cell length, measured from the cell pole. For McdB^PopTag^ condensates, the distribution was quantified by calculating the fractional cross-sectional area of the ribosome-excluded condensate region. The cross-section area of the condensate region and the cell was measured using the freehand selection tool in Fiji (*14*), and the fractional cross-section area was calculated as the ratio of condensate area to the total cell cross-section area. The quantified data were plotted and analyzed using GraphPad Prism version 11.0.0 (GraphPad Software, San Diego, CA). To calculate carboxysome volume and surface area, the segmented carboxysome surfaces were imported into UCSF ChimeraX and the Measure Volume and Area tool was used to determine volume and surface area for each carboxysome. The measured values were plotted using GraphPad Prism version 11.0.0. The volume and surface area values are not normalized to the bacterial cell size.

### Image analysis

#### Cell segmentation

Cell segmentation was performed using the Omnipose package in Python as previously (*2*). For segmentation, a Gaussian blur (σ = 0.066 μm) was applied to phase-contrast images prior to processing with a custom-trained model (*2*). Cells touching the image borders were excluded. Erroneous segmentations were manually corrected with the Omnipose GUI or omitted from analysis.

#### Focus count analysis

Images from multiple fields of view were acquired as described above and analyzed using a custom Python script for fluorescent focus detection and analysis (*2*). Incorrectly identified fluorescent foci were manually filtered from further analysis. Detection parameters were optimized for each compartment type (see Supplementary Table 5). The number of cells with 0, 1, 2, or ≥3 foci were quantified and visualized using GraphPad Prism (GraphPad Software, San Diego, CA).

#### Fluorescence intensity profile analysis

Fluorescence intensity profiles were generated using ImageJ (*15*) . For each cell, a straight line was drawn across the longitudinal axis. The fluorescence intensity along the line was recorded. Cell length was normalized by rescaling the profile coordinates from 0 to 1. Cells without McdA were grouped and analyzed separately based on the number of observed foci (1 or 2). Cells producing McdA were analyzed and plotted as a separate group. Intensity profiles were visualized using GraphPad Prism (GraphPad Software, San Diego, CA).

#### Kymograph analysis

Drift-corrected time-lapse image stacks were analyzed to generate kymographs representing fluorescence dynamics along the cell length. In Fiji (ImageJ) (*15*) (Schneider et al. 2012), a straight line was manually drawn along the long axis of each cell. The Multi-Kymograph tool was used to extract intensity profiles. For visualization, intensity values at each time point were normalized to the minimum and maximum intensity within that frame. The resulting kymographs display the temporal evolution of fluorescence distribution along the cell axis.

### Statistical Analysis

The experimental design, sample sizes, number of biological replicates, and statistical tests are summarized in Supplementary Table 4. In summary, distributions are presented as scattered plots, thick bar indicate the interquartile range around the median, and error bars represent a interquartile range. For XY graphs, shaded areas indicate the 95% CI. When applicable, P-values and statistical significance were evaluated using Kolmogorov-Smirnov (for distributions with at least 50 data points that do not assume normality) or One-Way ANOVA (for comparisons between two groups, with three data points each).

A biological replicate was defined as an independent transformation, culture, and induction performed on a different day, starting from a separate colony.

---

**Supplementary Figure 1.**
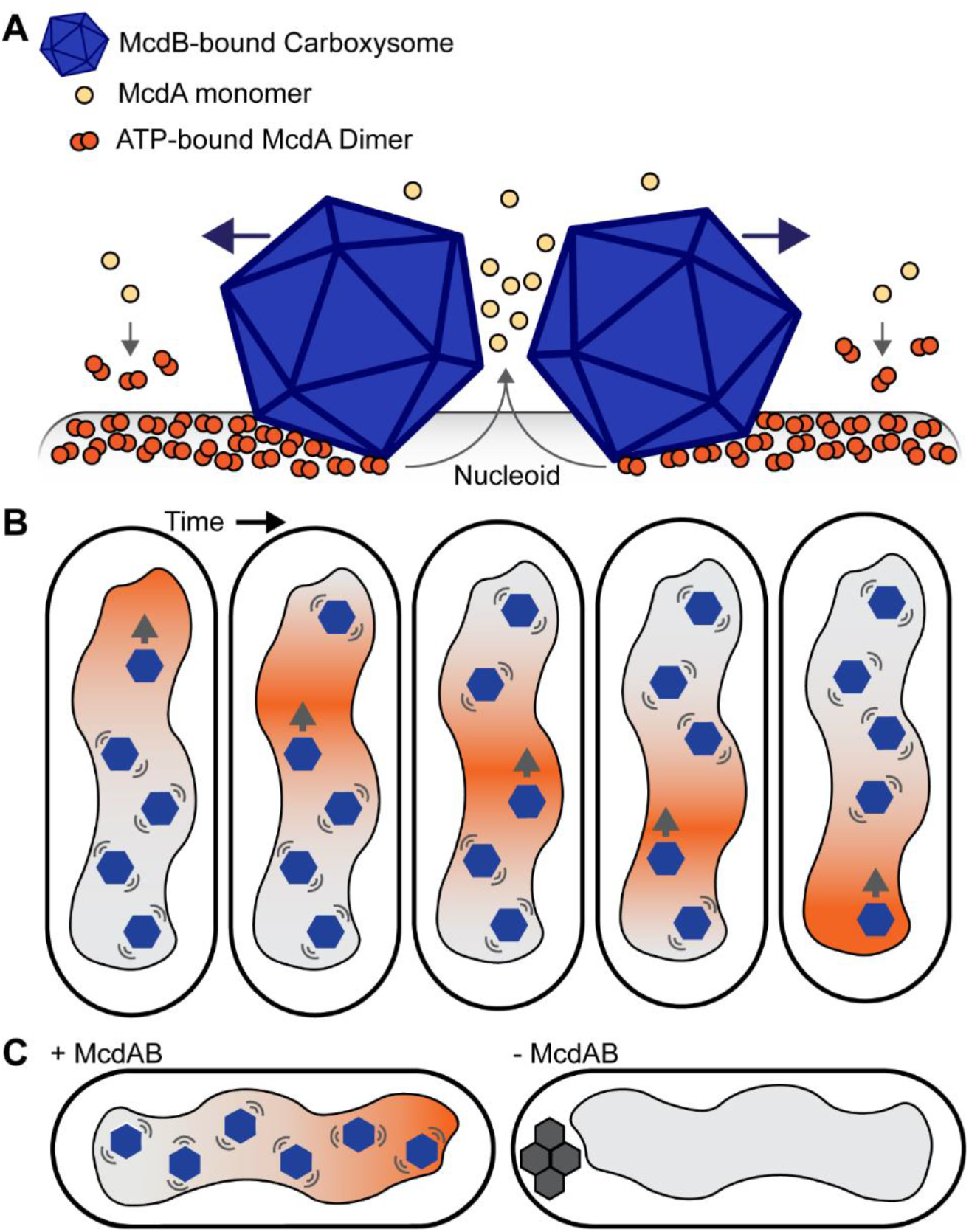
Carboxysomes are spatially organized by the McdAB system. (**A**) Diffusion-rachet model for carboxysome distribution over the nucleoid. McdB associates with the carboxysome shell. ATP-bound McdA dimerizes and binds the nucleoid via nonspecific DNA interactions. McdB stimulates McdA release from the nucleoid in the vicinity of McdB-bound carboxysomes. (**B**) The interaction between McdB-bound carboxysomes and nucleoid-localized McdA causes *(i)* an McdA depletion zone to form on the nucleoid in the vicinity of carboxysomes, *(ii)* a global break in McdA symmetry along the nucleoid, *(iii)* a movement of carboxysomes toward increased McdA concentrations, and *(iv)* a pole-to-pole oscillation of McdA that emerges across space and time. (**C**) Without the McdAB system, carboxysomes are mispositioned as nucleoid-excluded aggregates.

**Supplementary Figure 2.**
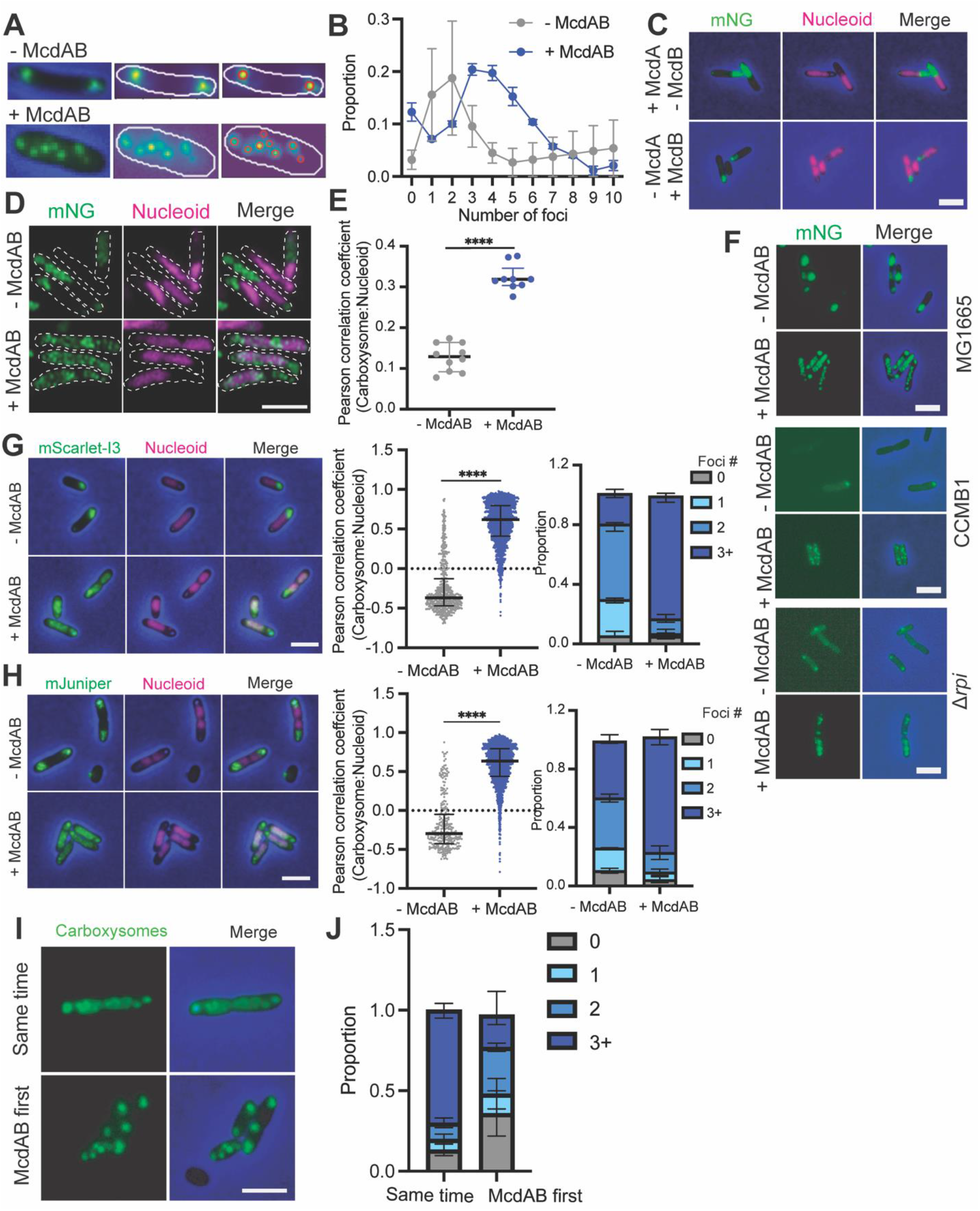
The McdAB system spatially organizes carboxysomes on the *E. coli* nucleoid. (**A**) Representative images of cells undergoing segmentation and focus detection pipeline. (**B**) Quantification of the proportion of cells with indicated numbers of carboxysome foci from Figure 1 A-B. (**C**) Wide-field fluorescence images of *E. coli* Top10 cells assembling carboxysomes in the presence of either McdA or McdB alone. mNG-labeled carboxysomes are shown in green, DAPI-stained nucleoids in magenta, and phase-contrast in blue; scale bar, 2 μm. (**D**) Super-resolution Airyscan images of *E. coli Top10* cells without (top) or with (bottom) the McdAB system, showing mNeonGreen-labeled carboxysomes (green), DAPI-stained nucleoids (magenta); merged images are shown at right. Dashed outlines indicate cell boundaries. Scale bar, 2 μm. (**E**) Pearson correlation coefficients measuring colocalization between mNeonGreen-labeled carboxysomes and DAPI-stained nucleoids from cells in (D). (**F**) Wide-field fluorescence images of *E. coli* cells with MG1655, CCMB1, and *Δrpi* mutant backgrounds assembling carboxysomes without or with the McdAB system. (**G**, **H**) Wide-field fluorescence images of cells assembling carboxysomes labeled with mScarlet-I3 (G) or mJuniper (H). Pearson correlation coefficients quantifying colocalization between labeled carboxysomes and DAPI-stained nucleoids, as well as the proportion of cells with 0, 1, 2, or ≥3 carboxysome foci, are provided. (**I**) Wide-field fluorescence images of *E. coli* Top10 cells assembling carboxysomes under conditions where McdAB is expressed simultaneously with carboxysomes (same time), or where McdAB is expressed prior to carboxysome induction (McdAB first). mNG-labeled carboxysomes are shown in green, DAPI-stained nucleoids in magenta, and phase-contrast in blue; scale bar, 2 μm. (**J**) Quantification of the proportion of cells in (I) exhibiting the indicated number of carboxysome foci.

**Supplementary Figure 3.**
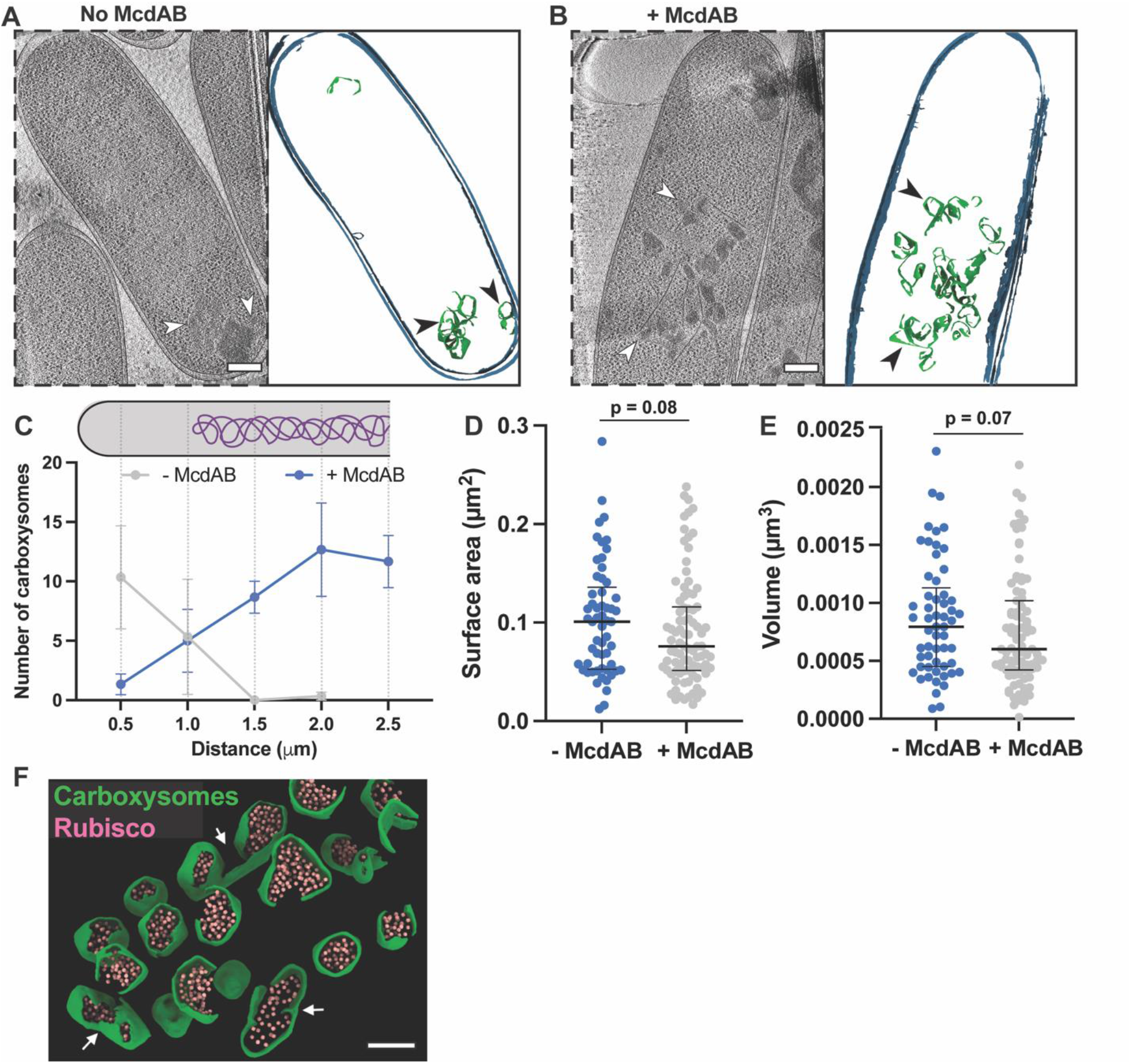
Analysis of carboxysomes positioning on the *E. coli* nucleoid. (**A, B**) Representative cryo-tomogram (left) and corresponding 3D segmentations (right) of *E. coli* cells showing carboxysome (arrowheads) distribution in the absence (A) or presence (B) of McdAB. For the segmentations, cell membranes are rendered in blue and carboxysomes in green. Scale bars, 100 nm. (**C**) Quantification of carboxysome number as a function of distance from the cell pole in cells from experiments shown in A and B. The schematic above the graph illustrates the positional bins measured along the cell’s long axis. (**D**, **E**) Comparison of individual carboxysome surface area (D) and volume (E) in cells from experiments shown in A and B. Each data point represents an individual measurement; bars represent mean ± SD; Statistical significance was determined by mann-Whitney test. (**F**) Zoomed-in view of a segmented tomogram showing carboxysome partitioning in *E. coli* expressing the McdAB system (same as Fig. 1N). The arrows mark the invaginations or partitions in the carboxysome shells. Scale bar: 100 nm.

**Supplementary Figure 4.**
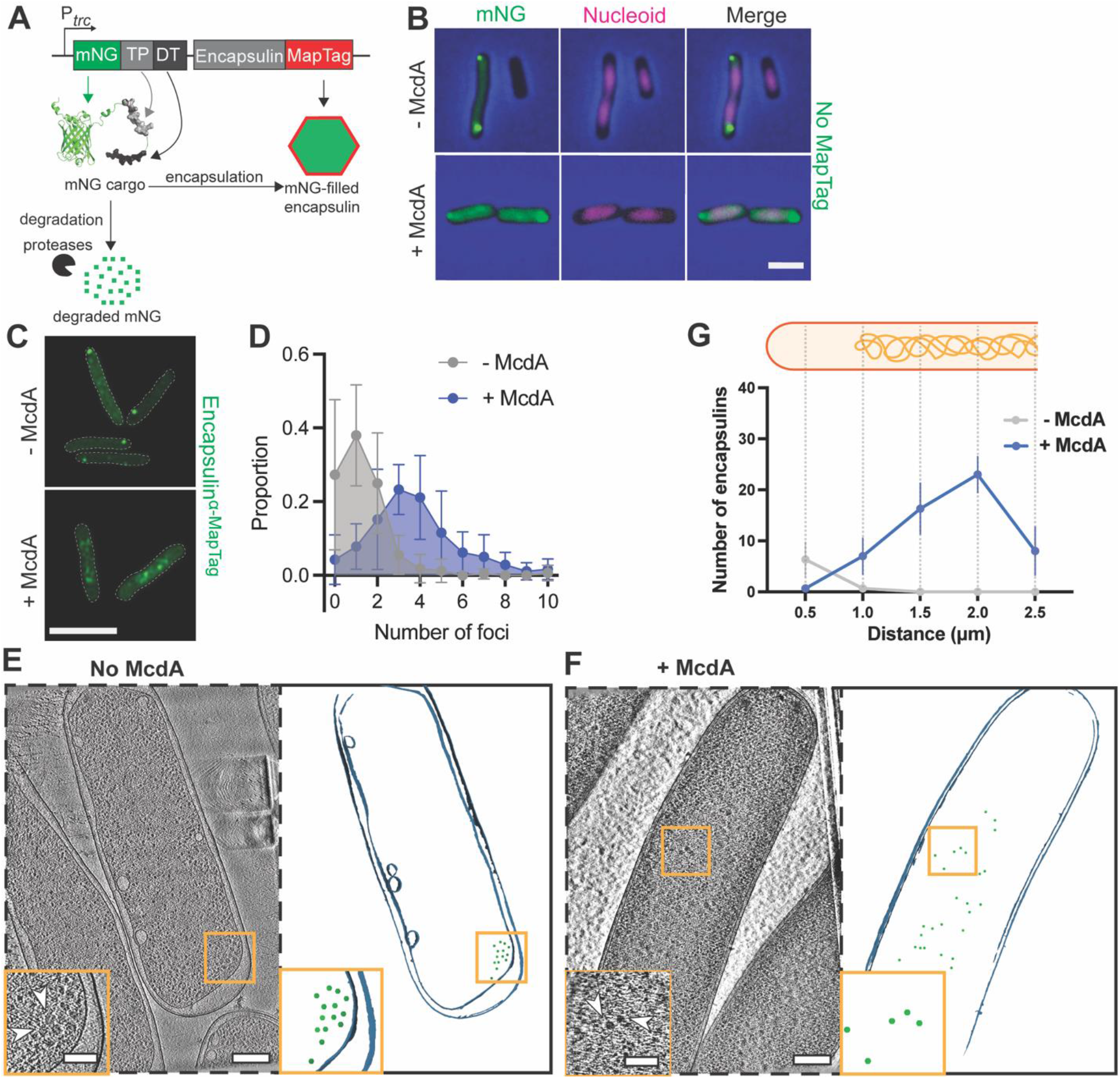
McdA spatially organizes both α- and β-MapTagged encapsulins on the *E. coli* nucleoid. (**A**) Schematic illustrating the visualization strategy for encapsulins. The cargo protein mNeonGreen (mNG) is fused to an encapsulin-specific targeting peptide (TP) and a degron tag (DT), while the encapsulin monomer is fused to the MapTag. Unencapsulated mNG is targeted for degradation by cellular proteases, thereby enhancing the contrast for visualization of mNG-filled encapsulins. (**B**) Wide-field fluorescence images of *E. coli* cells assembling encapsulins lacking the MapTag. Encapsulins are shown in green, DAPI-stained nucleoids in magenta, and phase-contrast in blue, in cells without (top) or with (bottom) McdA. Scale bar, 2 μm. (**C**) Super-resolution Airyscan images showing encapsulin^α-MapTag^ (green) in cells without (top) or with (bottom) McdA. Scale bars, 2 μm. (**D**) Proportion of cells with the indicated number of encapsulin^α-MapTag^ foci from (C). (**E, F**) Representative cryo-tomogram slices (left) and corresponding 3D segmentations (right) of *E. coli* cells showing encapsulin (arrowheads) distribution in the absence (E) or presence (F) of McdA. For the segmentations, cell membranes are rendered in blue and encapsulins in green. Scale bars, 100 nm. (**G**) Quantification of encapsulin number as a function of distance from the cell pole in cells from experiments shown in E and F. The schematic above the graph illustrates the positional bins measured along the cell’s long axis.

**Supplementary Figure 5.**
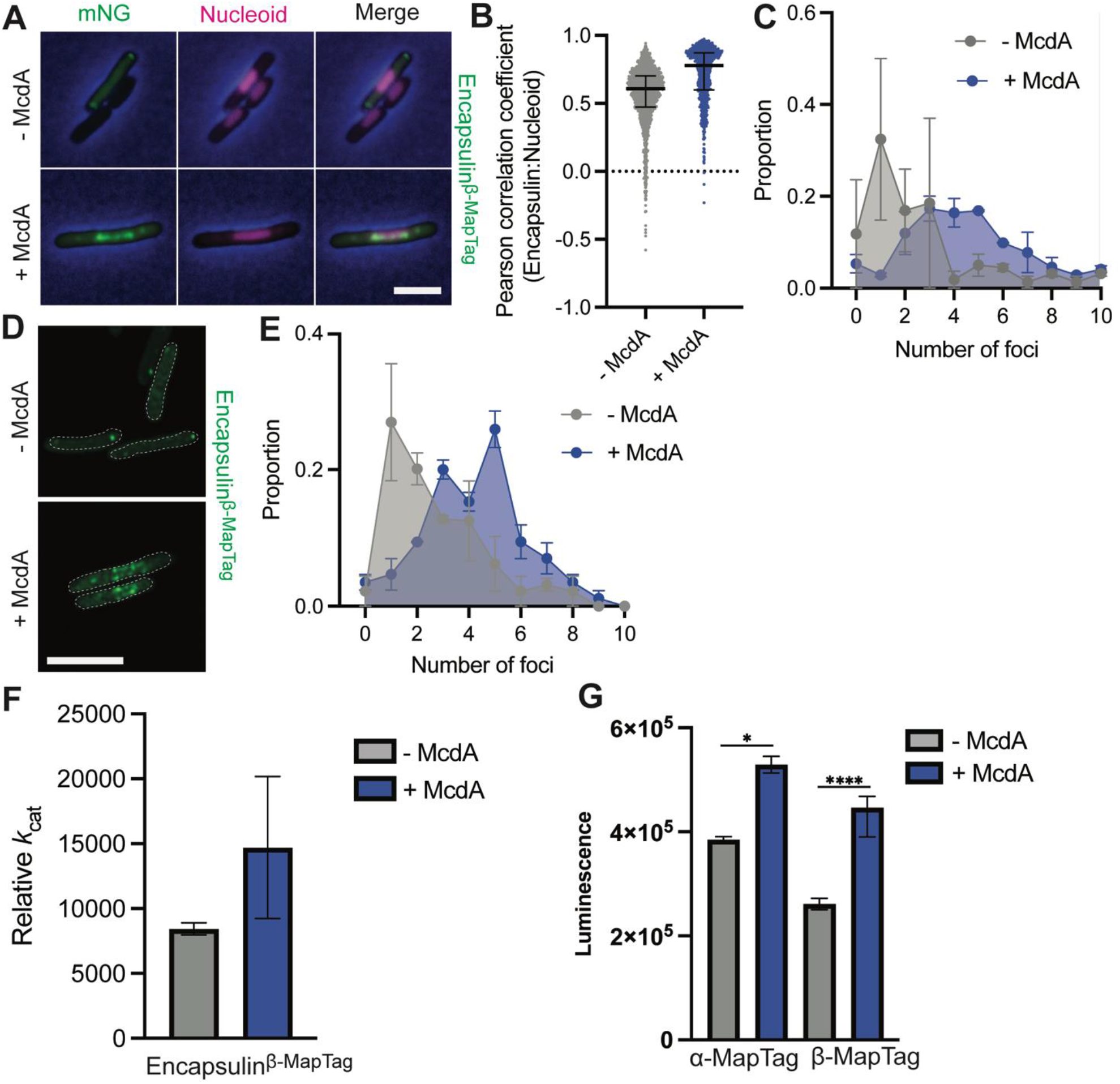
Positioning of encapsulins on the *E. coli* nucleoid and the resulting effects on encapsulated enzyme activity. (**A**) Wide-field fluorescence images showing encapsulin^β-MapTag^ (green), DAPI-stained nucleoids (magenta), and phase-contrast (blue) in cells without (top) or with (bottom) McdA. Scale bar, 2 μm. (**B**) Pearson correlation coefficients measuring colocalization between encapsulin^β-MapTag^ and nucleoids from (A). (**C**) Proportion of cells with the indicated number of encapsulin foci from (A). (**D**) Super-resolution Airyscan images showing encapsulin^β-MapTag^ (green) in cells without (top) or with (bottom) McdA. Scale bars, 2 μm. (**E**) Proportion of cells with the indicated number of encapsulin^β-MapTag^ foci from (D). (**F**) Relative *k*_cat_ of NanoLuc luciferase in cells containing mispositioned and positioned encapsulin^β-MapTag^. NanoLuc, instead of mNG, was targeted to encapsulins, and enzymatic activity was quantified by measuring luminescence following addition of the substrate furimazine. (**G**) Endpoint luminescence analysis of NanoLuc luciferase in cells containing mispositioned or positioned encapsulin^α-MapTag^ or encapsulin^β-MapTag^.

**Supplementary Figure 6.**
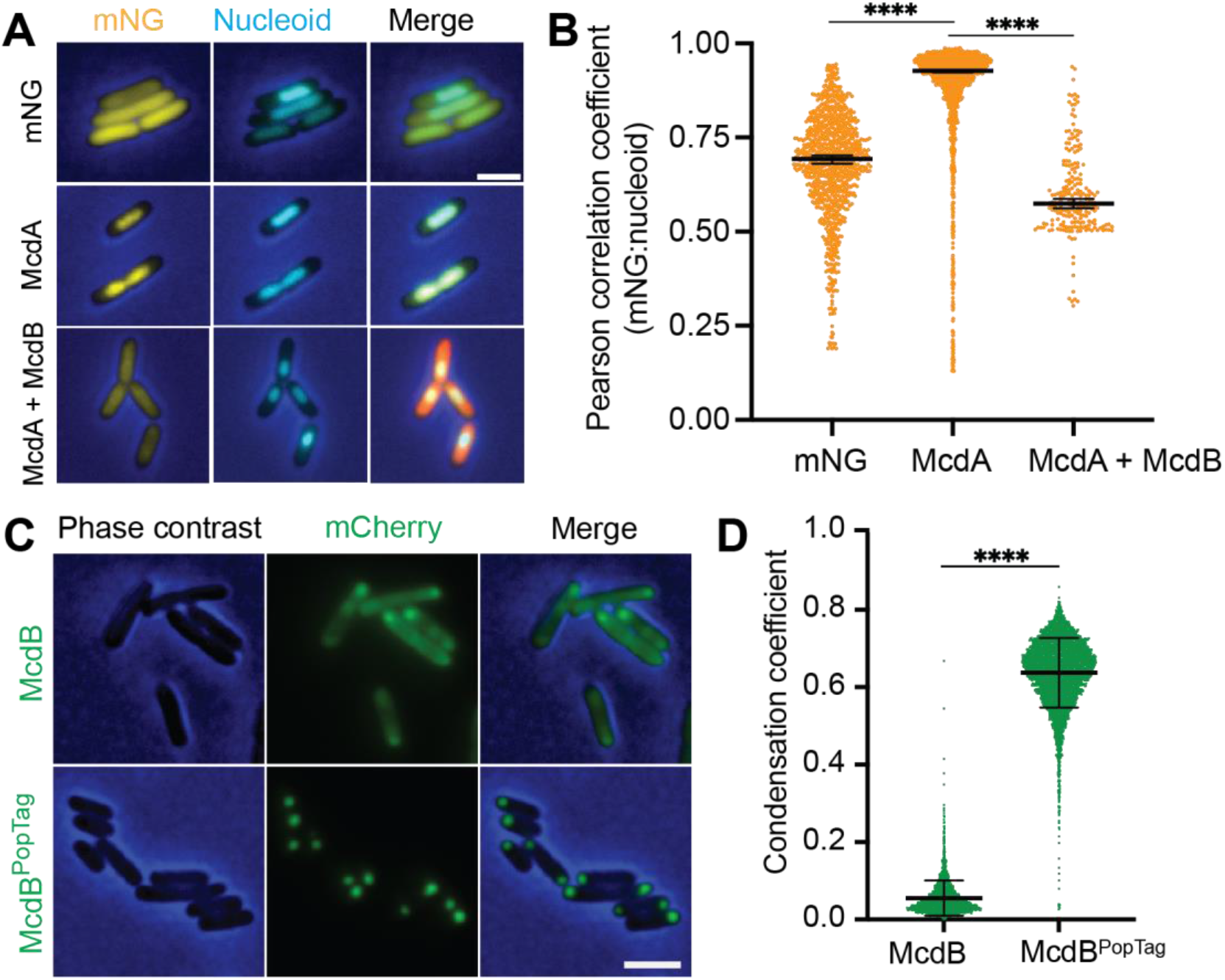
McdB globally releases McdA from the *E. coli* nucleoid. (**A**) Wide-field fluorescence images of cells expressing mNeonGreen (mNG), mNG-McdA, or both mNG-McdA and mCherry-McdB. Channels shown are mNG (yellow), DAPI-stained nucleoids (cyan), phase-contrast (blue), and mCherry (red), acquired 2 hours post-induction following chloramphenicol (Cm, 100 μg/mL) treatment to condense the nucleoid. Merged images are provided. Scale bar, 2 μm. (**B**) Pearson correlation coefficients measuring colocalization of mNG, mNG-McdA, or mNG-McdA in the presence of mCherry-McdB with the DAPI-stained nucleoid. (**C**) Wide-field fluorescence images showing McdB (top) or McdB^PopTag^ (bottom) condensates in *E. coli*. Phase-contrast (blue) and mCherry (green) are shown, with merged images. Scale bar, 2 μm. (**D**) Single-cell condensation coefficients of McdB and McdB^PopTag^ from (C).

**Supplementary Figure 7.**
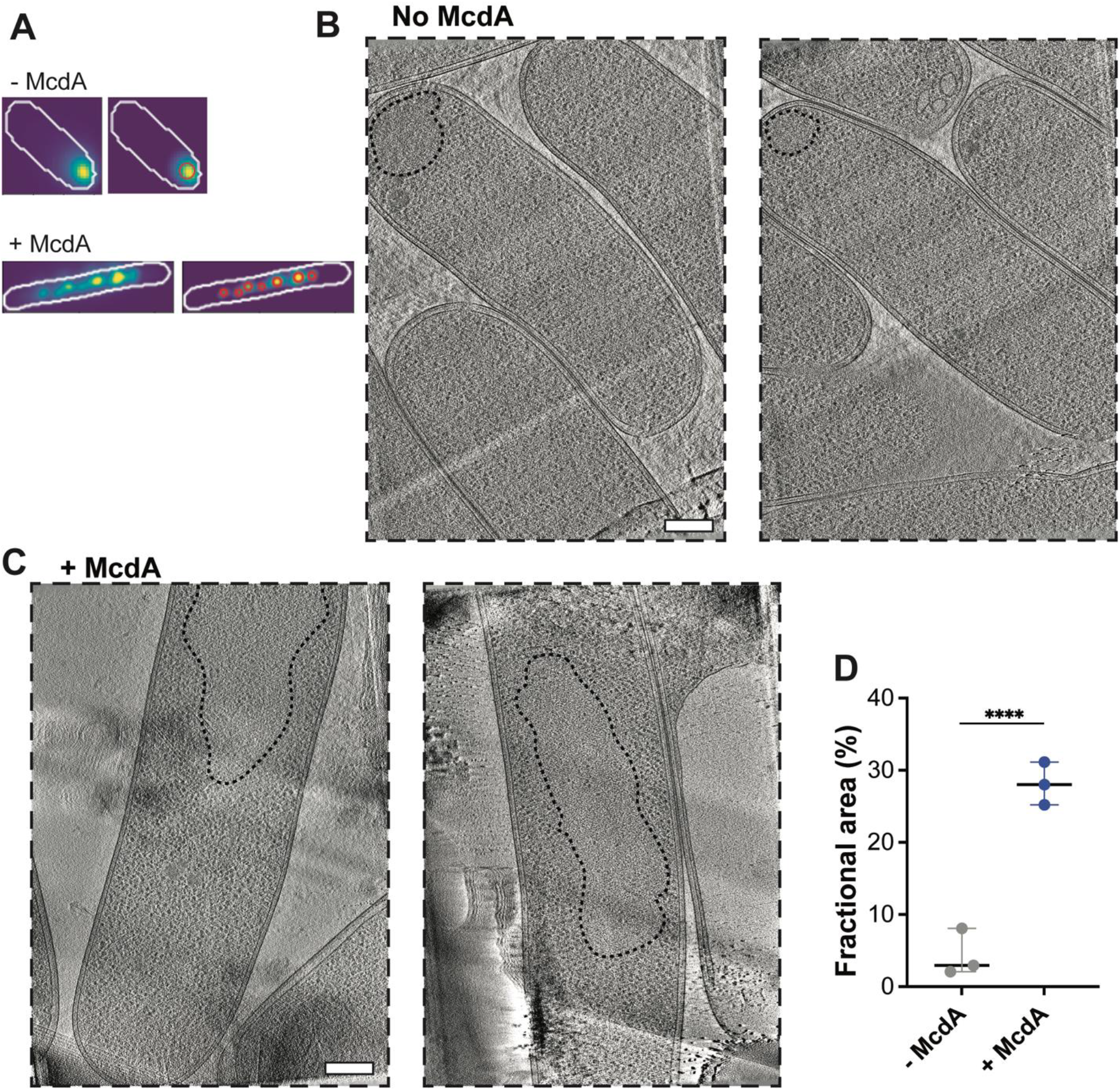
Positioning of condensates on the *E. coli* nucleoid. (**A**) Representative images of cells undergoing segmentation and focus detection pipeline. (**B**, **C**) Representative cryo-ET tomogram slices of *E. coli* cells showing condensate areas (dashed line) in the absence (B) or presence (C) of McdA. Scale bars, 100 nm. (**D**) Quantification of the fractional cell area occupied by condensates from experiments in B and C. Each data point represents an individual measurement; bars represent mean ± SD.

**Supplementary Figure 8.**
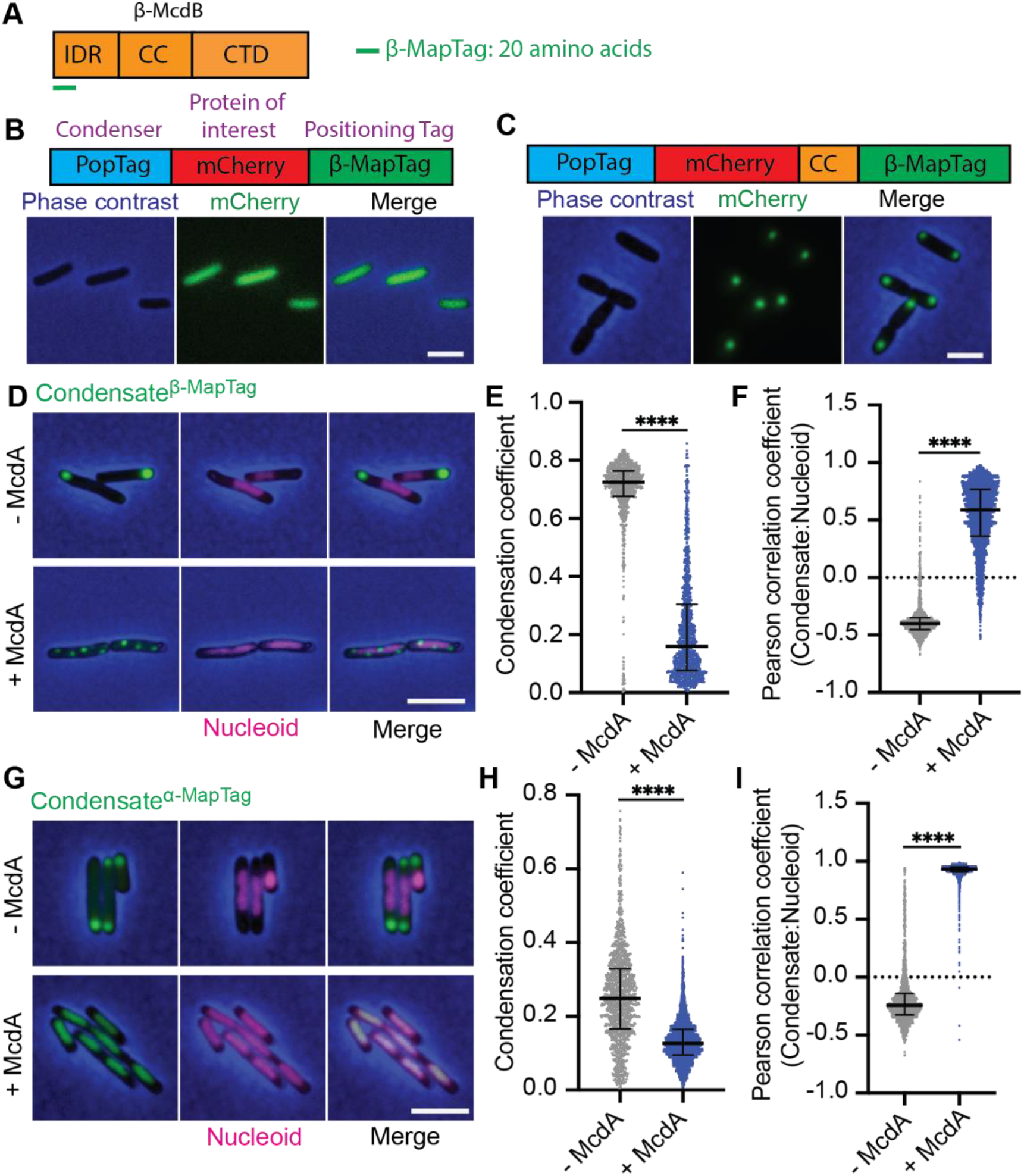
McdA spatially organizes both α- and β-MapTagged condensate monomers on the *E. coli* nucleoid. (**A**) Schematic of β-McdB (from *S. elongatus*) domain organization: intrinsically disordered region (IDR, including the β-MapTag - first 20 amino acids), coiled-coil (CC) region, and C-terminal domain (CTD). (**B**) Wide-field fluorescence images showing that mCherry^PopTag^ fused to β-MapTag does not form condensates. Phase-contrast (blue), mCherry (green), and merged images are shown. Scale bar, 2 μm. (**C**) Addition of the CC region restores cytoplasmic condensate formation to mCherry^PopTag^. (**D**, **G**) Wide-field fluorescence images of condensates fused to the β-MapTag (D) or α-MapTag (G), with condensates in green, DAPI-stained nucleoids in magenta, and phase-contrast in blue. Scale bars, 2 μm. (**E**, **H**) Single-cell condensation coefficients of proteins of interest fused to β-MapTag (E) or α-MapTag (H). (**F**, **I**) Pearson correlation coefficients for colocalization between condensates fused to β-MapTag (F) or α-MapTag (I) and DAPI-stained nucleoids.

**Supplementary Figure 9.**
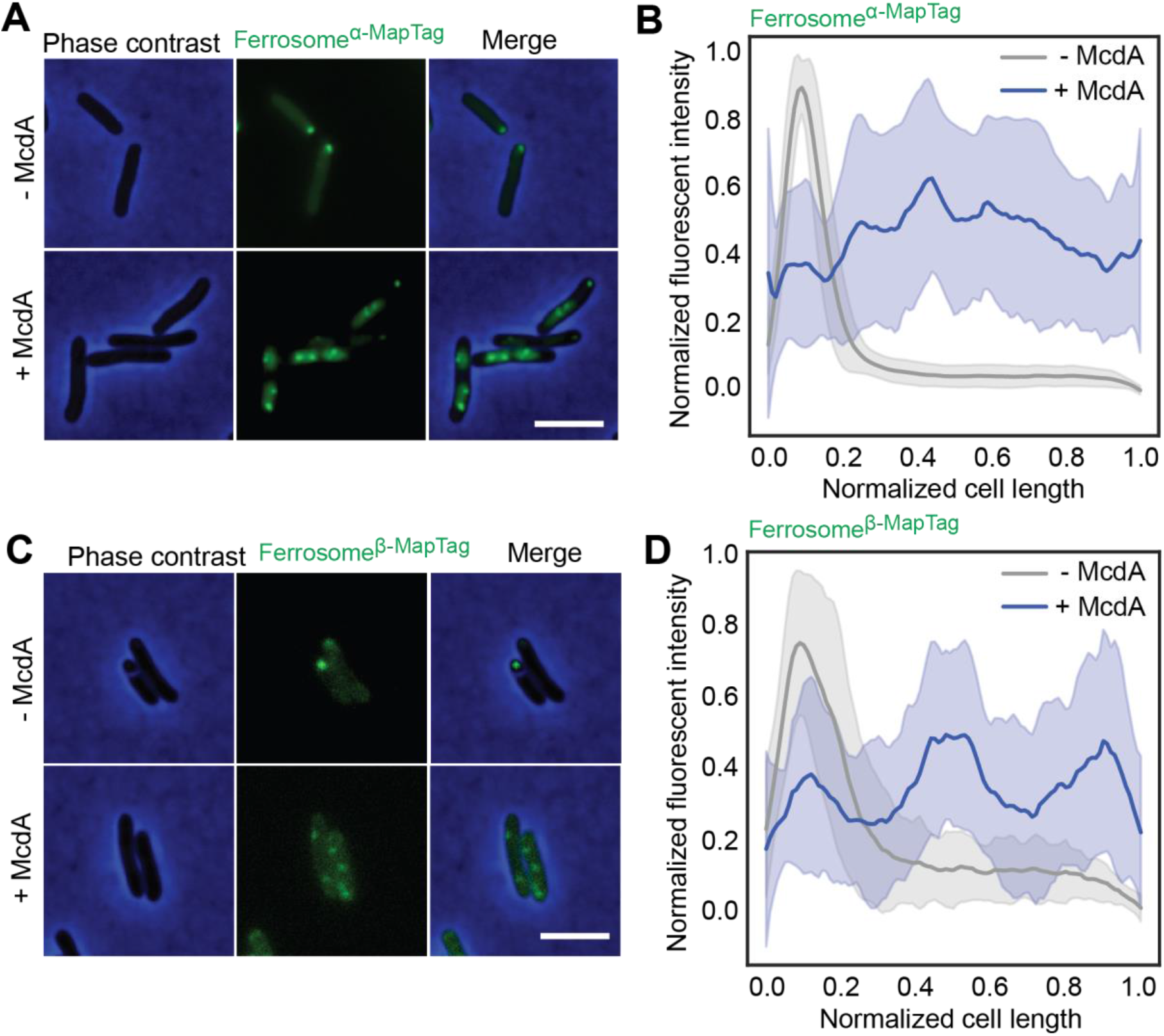
McdA spatially organizes both α- and β-MapTagged ferrosomes on the *E. coli* nucleoid. (**A**, **C**) Wide-field fluorescence images of ferrosomes fused to the α-MapTag (A) or β-MapTag (C), with ferrosomes in green and phase-contrast in blue. Scale bars, 2 μm. (**B**, **D**) Fluorescence intensity profiles of ferrosome signals measured along the normalized cell length in cells containing ferrosome^α-MapTag^ (B) or ferrosome^β-MapTag^ (D) foci.

**Supplementary Figure 10.**
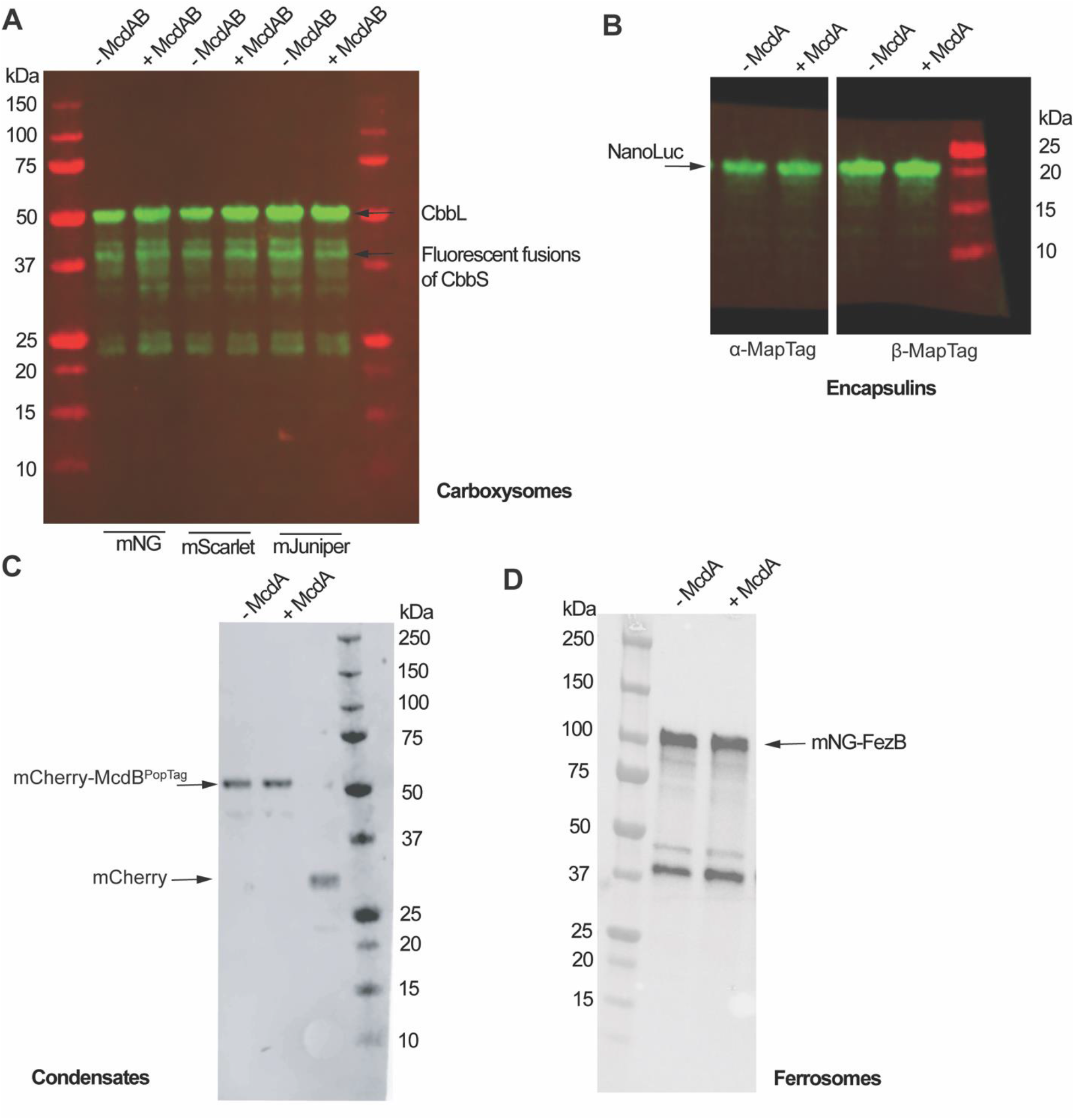
Immunoblots of fusion proteins. (**A**) Immunoblot of CbbS probed with anti-RbcL antibody. *E. coli* Top10 cells expressing the Rubisco small subunit CbbS fused to mNeonGreen (mNG), mScarlet, or mJuniper, in the presence or absence of McdAB. Arrows indicate the CbbL band and fluorescently tagged CbbS fusion proteins. (**B**) Immunoblot of NanoLuc luciferase probed with anti-NanoLuc antibody. *E. coli* MG1655 cells producing encapsulin^α–MapTag^ or encapsulin^β–MapTag^, in the absence or presence of McdA. NanoLuc luciferase was targeted to encapsulins. Arrow indicates the NanoLuc band. (**C**) Immunoblot of mCherry–McdB^PopTag^ probed with anti-mCherry antibody. *E. coli* MG1655 cells producing mCherry–McdB^PopTag^ in the absence or presence of McdA. Cells expressing mCherry alone were included as a control. Arrows indicate mCherry–McdB^PopTag^ or mCherry. (**D**) Immunoblot of mNG–FezB probed with anti-FezB antibody. *E. coli* MG1655 cells expressing mNG–FezB in the absence or presence of McdA. Arrow indicates the mNG–FezB band.

**Supplementary Table 1.** Strains and plasmids used in this study.

| Strains & Plasmids | Description | Source |
| --- | --- | --- |
| **Strains** |  |  |
| *E. coli* Top10 | F<sup>-</sup> mcrA $\Delta$(mrr-hsdRMS-mcrBC) ϕ80lacZΔM15 ΔlacX74 recA1 araD139 $\Delta$(ara-leu)7697 galU galK rpsL(StrR) endA1 nupG | Thermo Fisher Scientific |
| *E. coli* MG1655 ΔlacY | MG1655 derivative with a deletion of the lacY gene (lactose permease). | (16) |
| *E. coli* DH5alpha | F<sup>-</sup> ϕ80lacZΔM15 $\Delta$(lacZYA-argF)U169 recA1 endA1 hsdR17(rK<sup>-</sup> mK<sup>+</sup>) phoA supE44 λ- thi-1 gyrA96 relA1 | Thermo Fisher Scientific |
| **Plasmids** |  |  |
| pDT074 | Carboxysomes with CbbS C-terminally fused to mNeonGreen, Cm<sup>R</sup> | (1) |
| pDT081 | pDT074 + McdAB, Cm<sup>R</sup> | This study |
| pDT075 | Carboxysomes with CbbS C-terminally fused to mScarlet-I3, Cm<sup>R</sup> | (1) |
| pDT082 | pDT155 + McdAB, Cm<sup>R</sup> | This study |
| pDT078 | Carboxysomes with CbbS C-terminally fused to mJuniper, Cm<sup>R</sup> | (1) |
| pDT084 | pDT163 + McdAB, Cm<sup>R</sup> | This study |
| pSMK1 | Encapsulins with α-MapTag, Amp<sup>R</sup> | This study |
| pYH84 | P<sub>tet</sub>- mcdA with mcdA from *H. neapolitanus*, Km<sup>R</sup> | This study |
| pSMK2 | Encapsulins with β-MapTag, Amp<sup>R</sup> | This study |
| pYH91 | P<sub>tet</sub>- mcdA with mcdA from *S. elongatus*, Km<sup>R</sup> | This study |
| pRED1 | P<sub>tet</sub>-mNeonGreen, Km<sup>R</sup> | This study |
| pYH81 | P<sub>tet</sub>-mNeonGreen-mcdA with mcdA from *S. elongatus*, Km<sup>R</sup> | This study |
| pYH71 | P<sub>trc</sub>-mCherry-mcdB with mcdB from *S. elongatus*, Amp<sup>R</sup> | (2) |
| pYH82 | P<sub>trc</sub>-mCherry-poptag-mcdB with mcdB from *S. elongatus*, Amp<sup>R</sup> | This study |
| pJAB30 | P<sub>trc</sub>-mCherry-poptag-β-maptag, Amp<sup>R</sup> | This study |
| pJAB39 | pJAB30 with added coiled-coil region | This study |
| pRED2 | P<sub>trc</sub>-mCherry-poptag-α-maptag, Amp<sup>R</sup> | This study |
| pKM1 | Ferrosomes with α-MapTag, Amp<sup>R</sup> | This study |
| pKM2 | Ferrosomes with β-MapTag, Amp<sup>R</sup> | This study |

**Supplementary Table 2.**
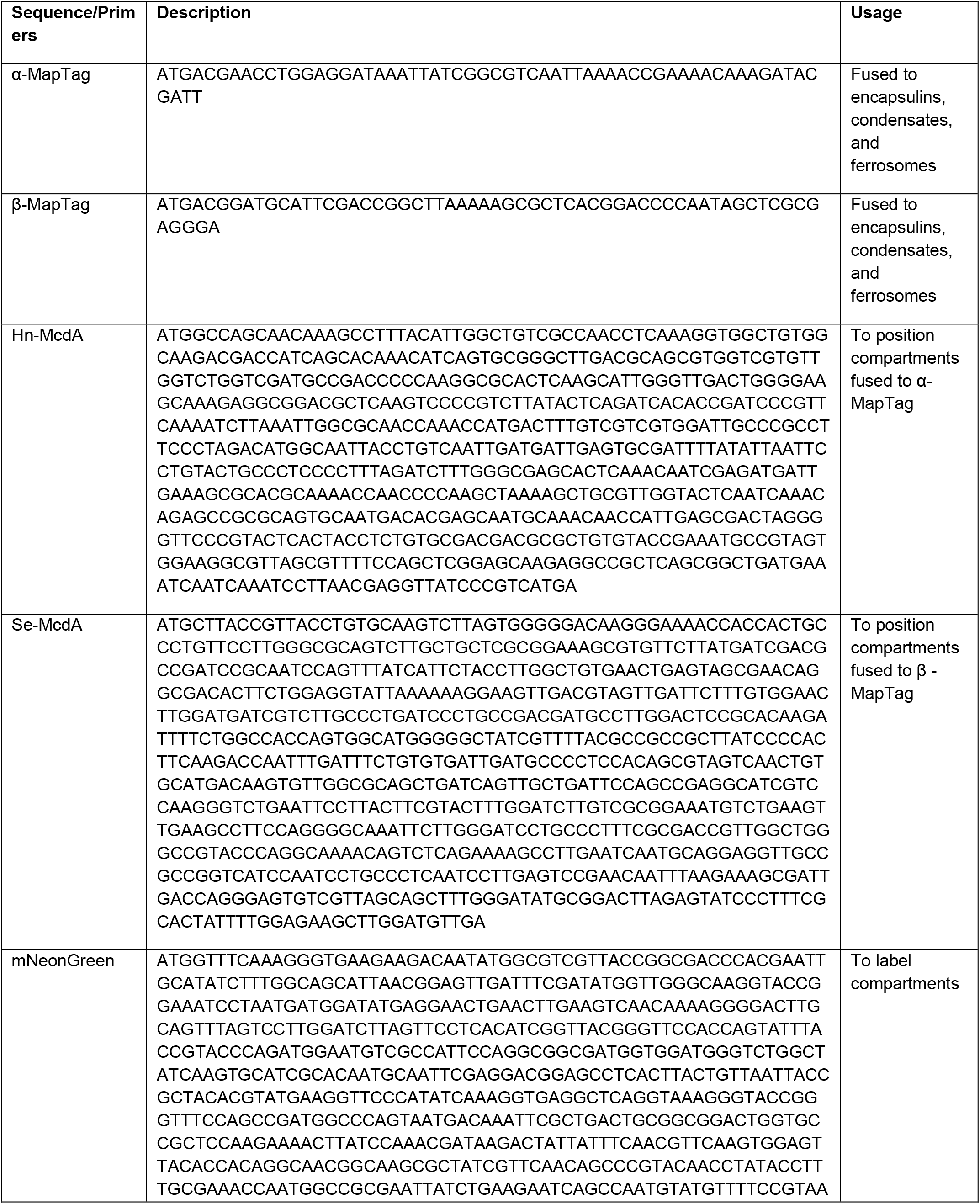
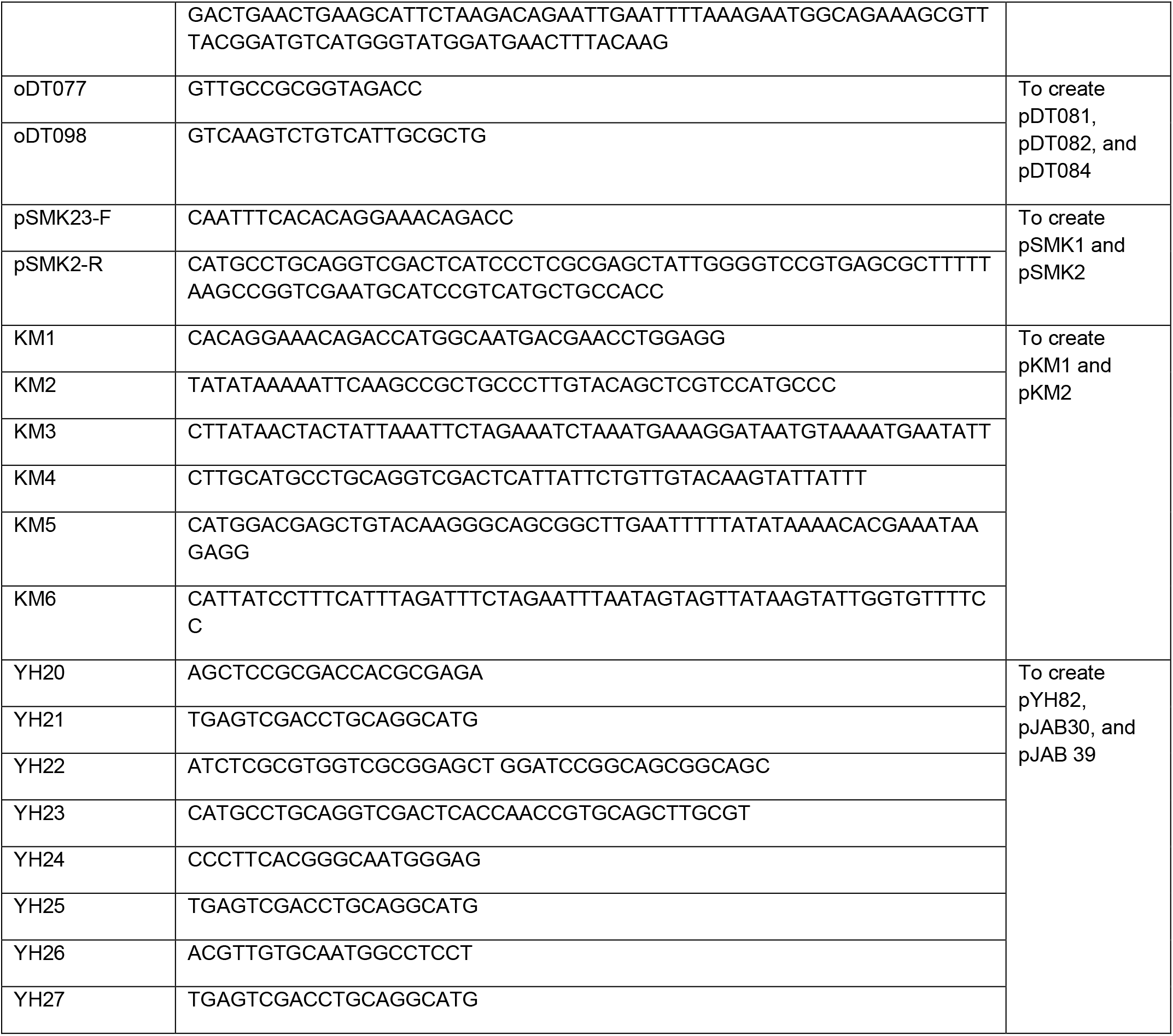
Synthetic DNA sequences & Primers used in this study.

**Supplementary Table 3.** Reagents used in this study.

| Reagent | Vendor | Catalog # | Use |
| --- | --- | --- | --- |
| Carbenicillin | Thermo Fisher Scientific | MT46100RG | To select for the plasmid in culture |
| Kanamycin | Sigma | K1377-5G |  |
| Chloramphenicol | Sigma | C0378-100G |  |
| DAPI | Thermo Fisher Scientific | D1306 | To label the nucleoid |
| Glucose | Thermo Fisher Scientific | 15023021 | As a carbon source and to inhibit protein production from the $P_{trc}$ promoter |
| Glycerol | Thermo Fisher Scientific | G334 | As a carbon source for *E. coli* MG1655 |
| IPTG | Thermo Fisher Scientific | 501126935 | To induce protein production |
| L-arabinose | Thermo Fisher Scientific | AC365181000 | To induce protein production |
| Anhydrotetracyclin |  |  |  |
| Casamino acids | Millipore Sigma | 65072-00-6 | For AB medium |
| Thiamine | Millipore Sigma | T4625-5G | For AB medium |
| Uracil | Millipore Sigma | 66-22-8 | For AB medium |
| UltraPure agarose | Invitrogen | 16500100 | For agar pads |
| KasI | New England Biolabs | R0544S | Restriction enzyme |
| BamHI-HF | New England Biolabs | R3136S | Restriction enzyme |
| NcoI-HF | New England Biolabs | R3193S | Restriction enzyme |
| Sall-HF | New England Biolabs | R3138S | Restriction enzyme |
| QIAquick Gel Extraction Kit | QIAGEN | Q28704 | Gel extraction |
| T4 DNA Ligase | New England Biolabs | M0202L | Cloning |
| NEBuilder HiFi DNA Assembly Master Mix | New England Biolabs | E2621L | Cloning |
| Phusion MM | New England Biolabs | M0531L | Cloning |
| QIAquick PCR Purification Kit | QIAGEN | 28106 | PCR purification |

**Supplementary Table 4.** Media and induction conditions.

| Figure | Growth medium | Induction conditions |  |
| --- | --- | --- | --- |
| 1A-B | LB | 0.2% arabinose | 2 h |
| 1E | LB | 0.2% arabinose | 3 h |
| 1H | LB | 0.2% arabinose | 5 h |
| 1 I | LB | 0.2% arabinose + 100 nM aTc for 2 h; cells were washed prior to 8 h of imaging. |  |
| 1 K | LB | 0.2% arabinose for 2 h; 100 nM aTc was then added, and cells were imaged for 6 h. |  |
| 1 M-N | LB | 0.2% arabinose | 2 h |
| 2A | AB | 1 mM IPTG | 2 h |
| 2B | AB | 1 mM IPTG; 20 nM aTc | 2h |
| 2E | AB | 1 mM IPTG | 6 h |
| 2G | AB | 1 mM IPTG; 20 nM aTc | 4 h |
| 2I | AB | 1 mM IPTG + 20 nM aTc for 2 h; cells were washed prior to 6 h of imaging. |  |
| 2K | AB | 1 mM IPTG for 2 h; 20 nM aTc was then added, and cells were imaged for 1 h. |  |
| 2M | AB | 1 mM IPTG | 2 h |
| 2N | AB | 1 mM IPTG; 20 nM aTc | 2 h |
| 3A | AB | 100 μM IPTG | 2 h |
| 3B | AB | 100 μM IPTG; 5 nM aTc | 2 h |
| 3E | AB | 100 μM IPTG | 3h |
| 3G | AB | 100 μM IPTG; 5 nM aTc | 6 h |
| 3I | AB | 100 μM IPTG + 5 nM aTc for 2 h; cells were washed prior to 6 h of imaging. | |
| 3K | AB | 1 mM IPTG for 2 h; 20 nM aTc was then added, and cells were imaged for 5 h. |  |
| 3M | AB | 100 μM IPTG | 2 h |
| 3N | AB | 100 μM IPTG; 5 nM aTc | 2 h |
| 4A | AB | 1 mM IPTG | 2 h |
| 4B | AB | 1 mM IPTG; 20 nM aTc | 2 h |
| 4E | AB | 1 mM IPTG; 20 nM aTc | 3 h |
| 4G | AB | 1 mM IPTG for 2 h; 20 nM aTc was then added, and cells were imaged for 3 h. |  |
| 4I | AB | 1 mM IPTG | 2 h |
| 4J | AB | 1 mM IPTG; 20 nM aTc | 2 h |
| S2C-H | LB | 0.2% arabinose; 100 nM aTc | 2 h |
| S3A-E | LB | 0.2% arabinose; 100 nM aTc | 2 h |
| S4B-H | AB | 1 mM IPTG; 20 nM aTc | 2 h |
| S5A-B | AB | 1 mM IPTG; 20 nM aTc | 2 h |
| 6A | AB | 100 μM IPTG; 5 nM aTc | 2 h |
| 6C | AB | 100 μM IPTG | 2 h |
| S7A | AB | 100 μM IPTG | 2 h |
| S7B | AB | 100 μM IPTG; 5 nM aTc | 2 h |
| S8B-C | AB | 1 mM IPTG | 2 h |
| S8D-G | AB | 100 μM IPTG; 5 nM aTc | 2 h |
| S9A-C | AB | 100 μM IPTG; 5 nM aTc | 2 h |
| S10A | LB | 0.2% arabinose; 100 nM aTc | 2 h |
| S10B | AB | 1 mM IPTG; 20 nM aTc | 2 h |
| S10C | AB | 100 μM IPTG; 5 nM aTc | 2 h |
| S10D | AB | 1 mM IPTG; 20 nM aTc | 2 h |

**Supplementary Table 5.** Statistical information from datasets in this study.

| Figure | Experiment | N cells (N replicates) | Statistical Test |
| --- | --- | --- | --- |
| 1C, D | Carboxysomes – mispositioned | 344 (3) | Kolmogorov-Smirnov |
|  | Carboxysomes – positioned | 1514 (3) |  |
| 1F | Carboxysomes – mispositioned over time | 3246 (3) |  |
| 1H | Carboxysomes – positioned over time | 8942 (3) |  |
| 2C | Encapsulins – mispositioned & positioned | 1759 (3) + 937 (3) | Kolmogorov-Smirnov |
| 2D | Encapsulins – mispositioned & positioned | 1759 (3) + 937 (3) |  |
| 2F | Encapsulins – mispositioned over time | 780 (3) |  |
| 2H | Encapsulins – positioned over time | 345 (3) |  |
| 2J | Encapsulins – McdA removal | 1080 (3) |  |
| 2L | Encapsulins – McdA addition | 369 (3) |  |
| 3C | Condensates – mispositioned & positioned | 20 (3) + 20 (3) |  |
| 3D | Condensates – mispositioned & positioned | 1423 (3) + 1872 (3) |  |
| 3J | Condensates – McdA removal | 17555 (3) |  |
| 3L | Condensates – McdA addition | 625 (3) |  |
| 4C | Ferrosomes – mispositioned & positioned | 179 (3) + 197 (3) | Kolmogorov-Smirnov |
| 4D | Ferrosomes – mispositioned & positioned | 179 (3) + 197 (3) |  |
| 4F | Ferrosomes – positioned over time | 3439 (3) |  |
| 4H | Ferrosomes – McdA addition | 2676 (3) |  |

**Supplementary Table 6.**
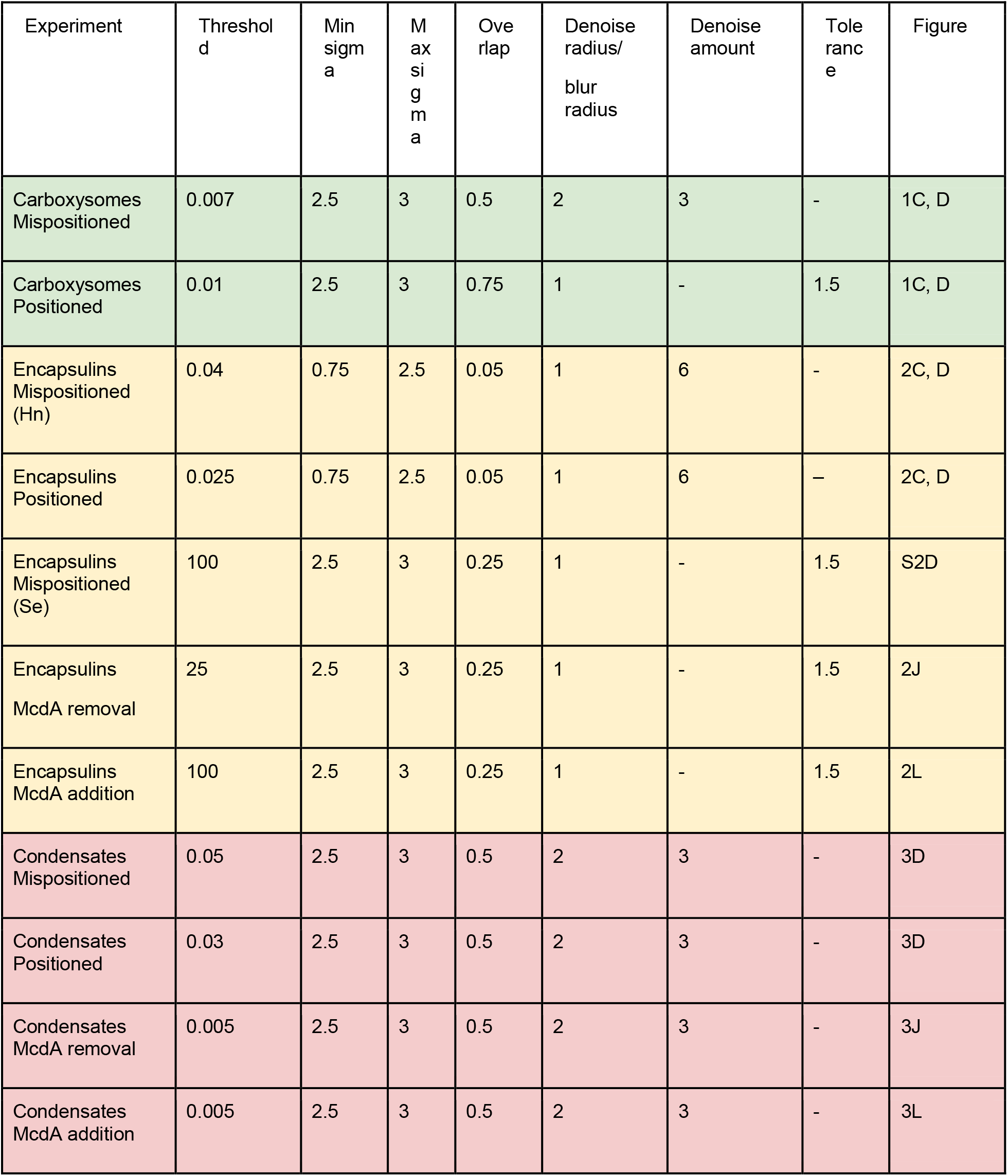
Detection parameters for image analysis.

| Experiment | Threshold | Min sigma | Max sigma | Overlap | Denoise radius/<br>blur radius | Denoise amount | Tolerance | Figure |
| --- | --- | --- | --- | --- | --- | --- | --- | --- |
| Carboxysomes Mispositioned | 0.007 | 2.5 | 3 | 0.5 | 2 | 3 | - | 1C, D |
| Carboxysomes Positioned | 0.01 | 2.5 | 3 | 0.75 | 1 | - | 1.5 | 1C, D |
| Encapsulins Mispositioned (Hn) | 0.04 | 0.75 | 2.5 | 0.05 | 1 | 6 | - | 2C, D |
| Encapsulins Positioned | 0.025 | 0.75 | 2.5 | 0.05 | 1 | 6 | - | 2C, D |
| Encapsulins Mispositioned (Se) | 100 | 2.5 | 3 | 0.25 | 1 | - | 1.5 | S2D |
| Encapsulins McdA removal | 25 | 2.5 | 3 | 0.25 | 1 | - | 1.5 | 2J |
| Encapsulins McdA addition | 100 | 2.5 | 3 | 0.25 | 1 | - | 1.5 | 2L |
| Condensates Mispositioned | 0.05 | 2.5 | 3 | 0.5 | 2 | 3 | - | 3D |
| Condensates Positioned | 0.03 | 2.5 | 3 | 0.5 | 2 | 3 | - | 3D |
| Condensates McdA removal | 0.005 | 2.5 | 3 | 0.5 | 2 | 3 | - | 3J |
| Condensates McdA addition | 0.005 | 2.5 | 3 | 0.5 | 2 | 3 | - | 3L |

**Video S1. The McdAB system spatially organizes carboxysomes on the *E. coli* nucleoid.** Time-lapse wide-field fluorescence microscopy of cells with (left) and without (right) the McdAB system. mNG-labeled carboxysomes are shown in green and phase-contrast in blue. Scale bar, 5 μm.

**Video S2. Reversible control of carboxysome positioning by McdAB manipulation.** Time-lapse wide-field fluorescence microscopy shows carboxysome positioning in response to McdAB removal (left) or addition (right). mNG-labeled carboxsyomes are shown in green and phase-contrast in blue. Scale bars, 2 μm.

**Video S3. The McdAB system regulates both assembly and positioning of carboxysomes in *E. coli*.** *In situ* cryo-electron tomography of *E. coli* cells without (left) or with (right) the McdAB system. Carboxysomes are shown in green, Rubisco in pink, ribosomes in yellow, and the cell membrane in blue.

**Video S4. McdA distributes MapTagged encapsulins on the *E. coli* nucleoid.** Time-lapse wide-field fluorescence microscopy of cells with (left) and without (right) McdA. mNG-labeled encapsulins are shown in green and phase-contrast in blue; scale bars, 5 μm.

**Video S5. Reversible control of encapsulin positioning by McdA manipulation.** Time-lapse wide-field fluorescence microscopy shows encapsulin positioning in response to McdA removal (left) or addition (right). mNG-labeled encapsulins are shown in green and phase-contrast in blue. Scale bars, 2 μm.

**Video S6. McdA distributes McdB^PopTag^ condensates on the *E. coli* nucleoid.** Time-lapse wide-field fluorescence microscopy of cells with (left) and without (right) McdA. Condensates are shown in green and phase-contrast in blue. Scale bars, 5 μm.

**Video S7. Reversible control of McdB^PopTag^ condensate positioning by McdA manipulation.** Time-lapse wide-field fluorescence microscopy shows encapsulin positioning in response to McdA removal (left) or addition (right). Condensates are shown in green and phase-contrast in blue. Scale bars, 5 μm.

